# Functional genomic analysis reveals mechanisms of epigenetic interference in SARS-CoV-1 and SARS-CoV-2

**DOI:** 10.64898/2026.01.28.702465

**Authors:** Amber R. Paulson, Vincent Montoya, Jeffrey B. Joy

**Author notes:** current address (Amber R. Paulson).

## Abstract

Small viral RNAs (svRNA) are key factors in host adaptation and virulence and are previously identified in SARS-CoV-2 and SARS-CoV-1. Little is known about svRNA conservation among high-risk coronaviruses (subgenus *Sarbecovirus*) or how svRNAs might contribute to COVID-19. Here, using functional genomics we identify six novel svRNA-triplex-forming oligonucleotides (svRNA-TFOs) in SARS-CoV-2, and through comparative analyses with SARS-CoV-1, delineate evolutionary pathways for svRNA-TFOs in *Sarbecovirus*. In addition to significant enrichment of the svRNA-TFO enhancer-associated gene targets among differentially expressed gene sets collected during SARS-CoV-2 infection, we find statistical support for non-random associations with >25 % of recombination breakpoints and hotspots in the Wuhan-Hu-1 genome. Small RNA-sequencing reveals highly abundant svRNA-TFO-N derived from the nucleocapsid of SARS-CoV-2. Using synonymous site conservation analysis, we find svRNA-TFO-N, and other svRNA-TFOs are conserved among both SARS-CoV-2- and SARS-CoV-1-related lineages. Furthermore, BLASTn reveals svRNA-TFO-N, svRNA-TFO-S.1 and svRNA-TFO-E share sequence homology against various mammalian genomes supporting a potential mechanism of host adaptation. Finally, we identify potential links between the putative svRNA-TFO-S.1 and svRNA-TFO-N RNA progenitor structures with variant of concern-associated S:D614G and N:D377Y mutations, respectively. Collectively these findings support a role for epigenetic interference with potential to reveal critical insights into the evolution of SARS-CoV-2 and SARS-CoV-1.

## INTRODUCTION

Small viral RNAs (svRNAs) are emerging as major players in the pathogenesis of viral infections [1–4], including in SARS-CoV-2, which produces both canonical microRNAs (miRNAs) and nuclear-acting microRNAs (namiRNAs) that regulate host gene expression at post-transcriptional and transcriptional levels, respectively [5–9]. More broadly, small non-coding RNAs (ncRNAs) less than 200 nt in length are well-established regulators of gene expression [10–12], with canonical ∼ 22 nt miRNAs being the most comprehensively studied; these single-stranded RNAs typically act through the RNA-induced Silencing Complex (RISC) to target the 3′ untranslated regions (UTRs) of messenger RNAs [13–16]. During SARS-CoV-2 infection, several virus-derived miRNAs, including those from the ORF7a and nsp2 regions, have been shown modulate host immune response [5,8] and correlate with disease severity, respectively [9]. In another study, an ORF8-encoded miRNA was shown to be abundant on nasal swabs collected from COVID-19 patients and suggested to act as a decoy by interacting with host miRNAs [6]. Additionally, SARS-CoV-2 also produces five human-identical sequence (HIS) namiRNAs that act through nuclear AGO2 to target human enhancers and upregulate adjacent or distal genes, including cytokine genes and *hyaluronan synthase 2* [7]. While Sarbecoviruses such as SARS-CoV-1 and SARS-CoV-2 have been intensively studied, the evolutionary processes underlying their emergence into the human population remains unresolved, underscoring the importance of understanding svRNA-mediated host interactions in this group.

While namiRNAs are generally known to target enhancers [17–19], increasing attention is now being directed toward the role of RNA triplex-forming oligonucleotides (TFOs), which can also target enhancers and other important regulatory regions in the genome [20–22]. Endogenous TFOs in humans can be encoded on long or small ncRNAs and range from ∼ 12–30 nt in length [23–32]. Their sequence-specific binding of polypurine DNA within the major groove is a result of three TFO motif types (GA, GU, or CU), which form RNA:DNA:DNA triple helices (triplexes) via Hoogsteen base-pairing [33–36]. Emerging *in silico*, biophysical, and *in vitro* evidence reveals a striking enrichment of TFO-binding sites within promoter regions, 5′ UTRs, super-enhancers, and repetitive elements [20,22]. One study showed that these sites are preferentially positioned within nucleosome-depleted regions and active regulatory elements where chromatin architecture promotes triplex formation [21]. These findings support a regulatory role for RNA-DNA heterotriplexes at key genomic loci. Despite the growing recognition of TFOs in eukaryotic gene regulation [37–39], their role in host–virus interactions remain largely unexplored. Host miRNAs, for instance, can form triplexes with integrated provirus genomes to regulate Lentivirus latency and persistence [40] and inhibit Human immunodeficiency virus 1 replication [41], and Influenza A virus can trigger endogenous triplex formation that represses transcription of the gene for beta-interferon [42]. Nevertheless, the potential contribution of exogenous virus-derived TFOs as a novel class of svRNA has not been investigated, making this study the first to explore this possibility.

Production of structural and accessory proteins in coronaviruses, involves a unique mechanism of discontinuous transcription and results in a nested set of subgenomic RNAs (sgRNAs) [43,44]. Transcriptome-wide studies using DNA Nanoball sequencing, direct Nanodrop sequencing, and small RNA-sequencing (RNA-seq) have revealed that SARS-CoV-2, beyond production of canonical sgRNAs, also generates a variety of non-canonical sgRNAs with putative RNA modifications at an AAGAA motif [43]. Additionally, three canonical miRNAs are produced by SARS-CoV-2, as mentioned [5,6,8]. Although less studied, the SARS-CoV-1 transcriptome also encodes three canonical miRNAs derived from nsp3 and N-ORF regions validated by Morales et al., one of which modulates lung pathology in mice [4]. Together, these findings highlight *Sarbecovirus* transcriptomes encode diverse RNA products with possible regulatory functions, yet it remains unclear whether such factors drive evolution or contribute to cross-species transmissibility among this high-risk group. To address this gap, we leverage genomic resources from diverse Sarbecoviruses alongside comparative small RNA-seq of SARS-CoV-2 and SARS-CoV-1 during Calu-3 cell infection [45].

We hypothesized that SARS-CoV-2 can produce svRNAs that are capable of triplex-mediated epigenetic interference thereby altering the expression of distal host genes relevant to infection. Epigenetic dysregulation is already suspected to be central to COVID-19, where distinct genome-wide DNA methylation profiles are detected in the blood of patients with severe COVID-19 [46,47]. Also, restructuring of the chromatin architecture by SARS-CoV-2 has been shown [48] and the virus produces nuclear-acting proteins, such as ORF8 that targets the host nucleosome by mimicking histone and disrupts epigenetic regulation [49]. Despite these clues, and the presence of viral namiRNAs in the nucleus [7], the role of triplex-forming svRNAs in SARS-CoV-2 has not been investigated, which motivates this study.

Importantly, previous studies validating canonical miRNAs in SARS-CoV-2 and SARS-CoV-1 [4,5,8,9] quantified their relative abundance using small RNA-seq library preparation methods that optimize for capture of canonical miRNAs with intact 5′ phosphate and 3′ hydroxyl groups [50–52]. However, because the RNase L pathway plays a central role during SARS-CoV-2 infection [53–56], we focused on whether svRNAs might instead arise from RNase-mediated degradation products. Using small RNA-seq data from an adapter ligation-free library preparation method [45] that enables better recovery of RNase-derived fragments [50] carrying 5′ hydroxyl groups and 2′ or 3′ phosphates [51,52,57], we identified six putative svRNA-TFOs of concern and investigated their conservation and evolution across SARS-CoV-2, SARS-CoV-1, and other Sarbecoviruses. We further assessed for enrichment of their enhancer-associated target genes among differentially expressed genes from human lung tissue and lung-derived cell lines infected with SARS-CoV-2, and whole blood from severe COVID-19 patients. We identified that two of the predicted svRNA-TFO progenitor structures show mutation-associated changes in Variants of Concern (VoCs), and all six svRNA-TFOs exhibited non-random associations with known genomic recombination breakpoints and hotspots in the Wuhan-Hu-1 genome. To support this analysis, a functional genomics framework: “**Epi**genetic **V**iral **I**nterference through **R**NA **T**riplex **Ex**ploration” (Epi-VIRTEX) was developed that combines machine-learning for viral miRNA prediction, direct RNA-seq data for observable SARS-CoV-2 sgRNAs [43], small RNA-seq from infected Calu-3 cells [45], and recombinant-aware phylogenetics across *Sarbecovirus* genomes isolated from humans, pangolins, and greater horseshoe bats.

## RESULTS

### Putative svRNA-TFOs map to SARS-CoV-2 recombination breakpoints and hotspots

The current study focused on identifying small viral RNA-encoding triplex-forming oligonucleotides (svRNA-TFOs) in SARS-CoV-2 and whether they are likely to engage in svRNA-mediated epigenetic interference via triplex-formation against enhancer regions in the host’s genome. TFOs are typically identified from sequences by implementing DNA:DNA:RNA base-pairing algorithms that use canonical triplex-formation rules based on *in vitro* experiments [38]. By using the *triplexator* algorithm [58], we identified 18 TFOs from SARS-CoV-2 Wuhan-Hu-1 reference genome based on searches for putative TFO target sites within consensus enhancer annotations for human lungs [59] (Supplemental Table S1). This approach was reasoned based on findings of Cetin *et al.* (2019) and Maldanado *et al.* (2019) that endogenous TFOs have strong affinity for enhancer regions [20,21] and that SARS-CoV-2 is a respiratory pathogen known to replicate in, and cause damages to, human lung tissues [60].

As a proof-of-concept, the Epi-VIRTEX pipeline considered 446 miRNAs that were predicted in this study from observable SARS-CoV-2 sub-genomic RNAs based on direct RNA-seq [43] and 331 additional miRNAs predicted from the Wuhan-Hu-1 genome in other studies [61–65] (Supplemental Table S2). Of the 18 putative TFOs in SARS-CoV-2 described above, 14 were either located adjacent to, or overlapping with, at least one predicted miRNA precursor sequence (Supplemental Table S1). Next these information were integrated with reads per million (RPM)-normalized small RNA-seq coverage to confirm infection-relevant signal (≥ 40 RPM) by 24 hpi within the 25-nt flanks in 6 out of 18 (33 %) of the TFOs (Figure 1A–F). Also, little to no level of background coverage was noted from the mock-infected and untreated samples for this dataset, which also demonstrated consistency between A and B replicates libraries, providing confidence in these data (see Supplemental Figures S1 through S5).

**Figure 1:**
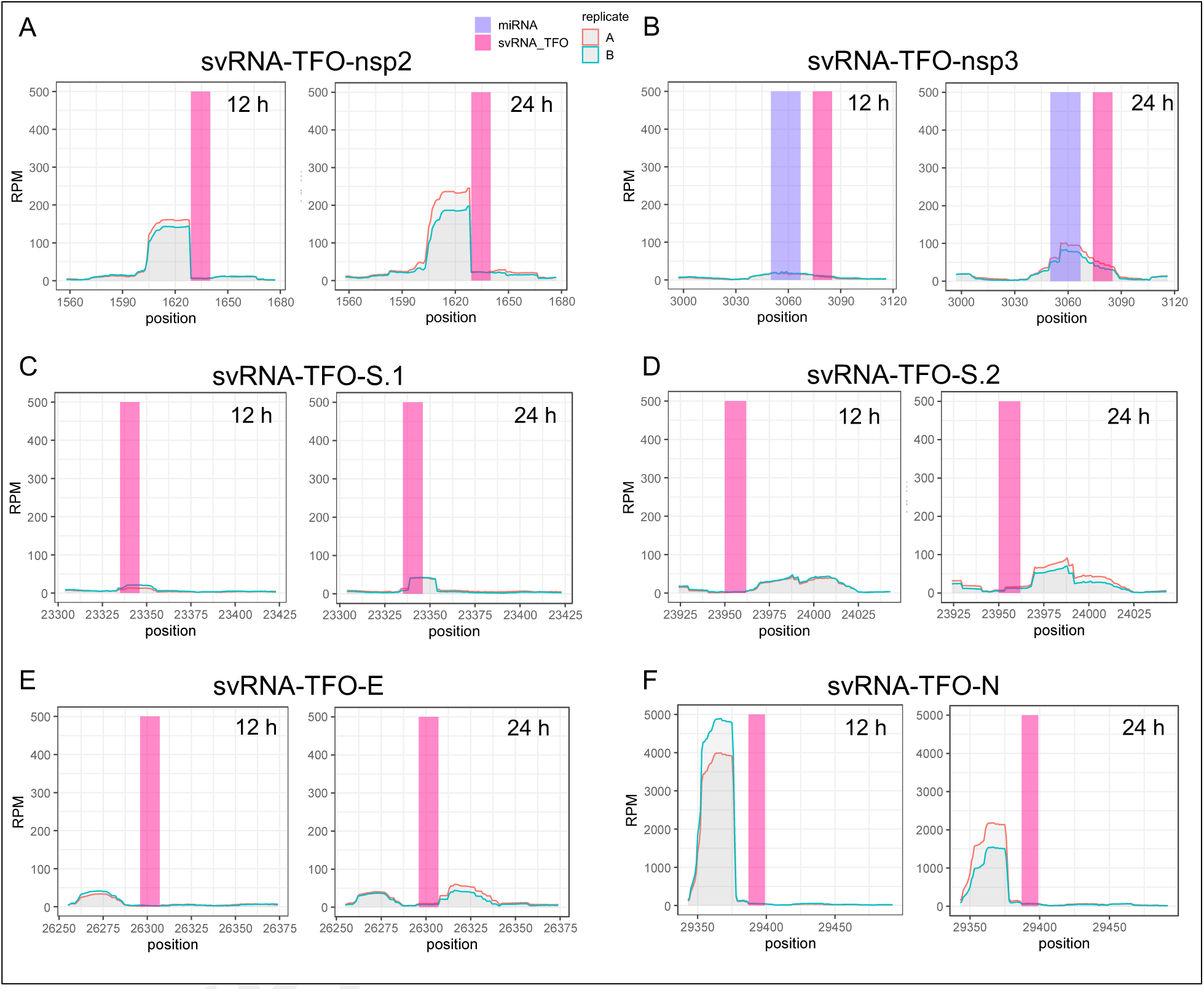
Small RNA-sequencing coverage across genomic regions encoding predicted SARS-CoV-2 small viral RNA (svRNA) triplex-forming oligonucleotides (TFOs) of concern. RPM-normalized small RNA-seq coverage is shown for A) svRNA-TFO-nsp2 (1,558 – 1,677); B) svRNA-TFO-nsp3 (2,997 –3,116) and svRNA-nsp3 (SARS-CoV-1) identified by Morales et al. [4]; C) svRNA-TFO-S.1 (23,304 – 23,423); D) svRNA-TFO-S.2 (23,924 – 24,043); E) svRNA-TFO-E (26,255 – 26,374); and F) svRNA-TFN-N (29,343 – 29,492) corresponding to the svRNA-TFO regions marked by numbered arrows and rectangles in Figure 2A. For the 4 h timepoints and controls (mock-infected and untreated) see Supplemental Figures S1 and S2).

These analyses suggest that SARS-CoV-2 produces six svRNAs-TFOs of concern, designated as: svRNA-TFO-nsp2, svRNA-TFO-nsp3, svRNA-TFO-S.1, svRNA-TFO-S.2, svRNA-TFO-E, and svRNA-TFO-N (Figure 2A). Importantly, we found all six of these svRNA-TFOs are located within close proximity (< 183 nt) of recombination breakpoints and hotspots previously identified in the Wuhan-Hu-1 reference genome [66] (Figure 2). A genome-wide analysis determined significant potential for co-localization between the svRNA-TFOs of concern and recombination breakpoints and hotspots (Wilcoxon adjusted [FDR] *p*-value = .0011). This was determined by comparing the average distance from the nearest recombination breakpoint or hotspot to the svRNA-TFOs of concern (49 nt ± 28 SEM) with the average distance from the remaining 12 TFOs (487 nt ± 85 SEM) and 1,200 randomly generated sites (710 nt ± 23 SEM) (Figure 2B). Using a similar but more conservative ORF-constrained approach, we observed significant statistical support (Wilcoxon adjusted [FDR] *p-*values < .05 to < .001) that svRNA-TFOs of concern are located closer to *bona fide* recombination breakpoints and hotspots rather than other TFOs and randomly generated sites (Figure 2C).

**Figure 2:**
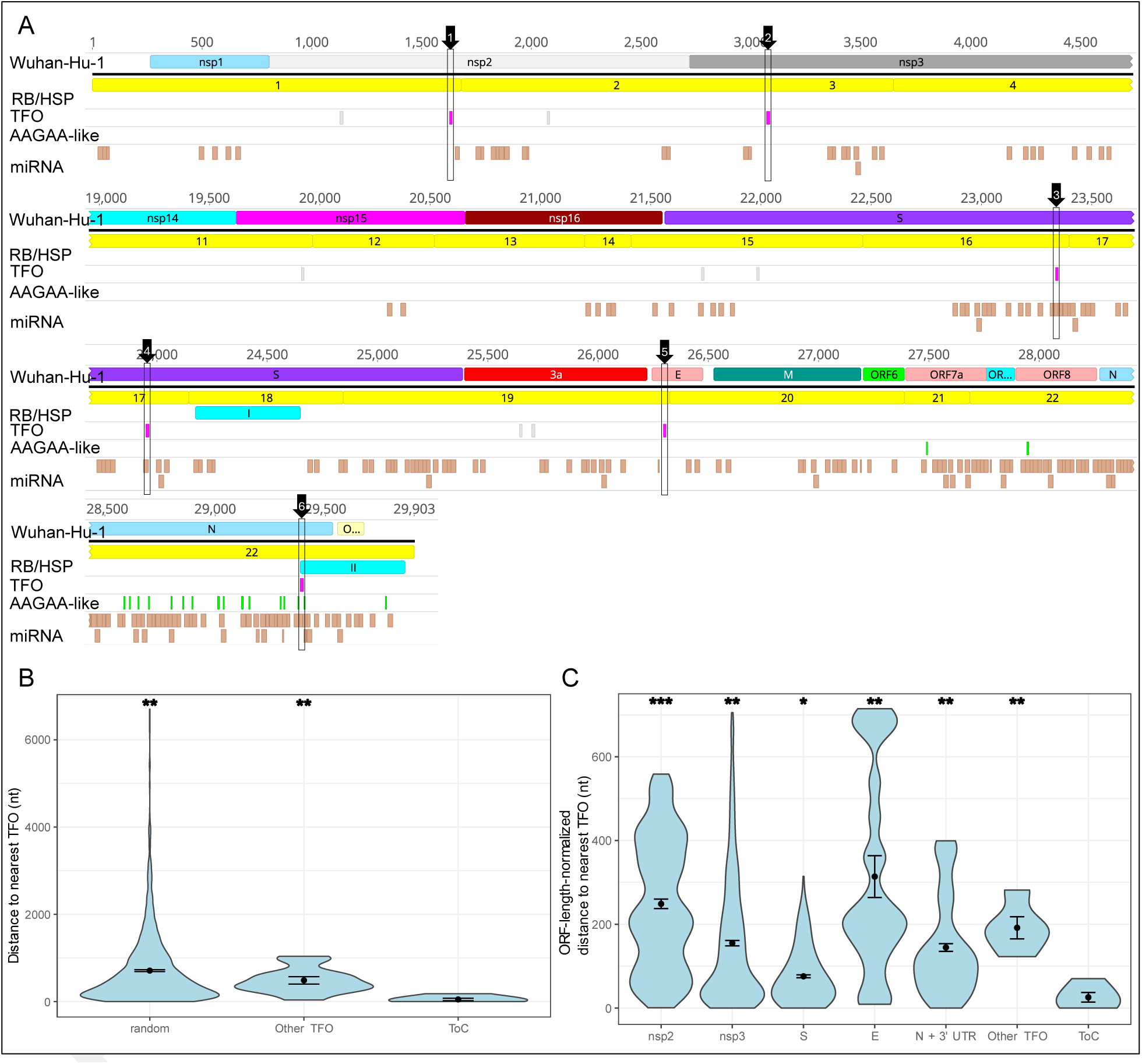
small viral RNAs encoding triplex-forming oligonucleotides (svRNA-TFOs) show non-random association with recombination break-points and hotspots in SARS-CoV-2. **A)** From top to bottom: **i)** Wuhan-Hu-1 reference genome (ASM985889) with main coding regions; **ii)** recombination breakpoint (RB) and hotspot (HSP) locations from Lytras *et al.* 2021 [66] shown from 1 – 22 in yellow, and I – II in cyan, respectively; **iii)** Triplex-forming oligonucleotides (TFOs) shown in light grey and pink with six svRNA-associated TFOs of concern (ToCs) indicated by numbered arrows and boxes (see also Figure 1 and Supplemental Table S1); **iv)** AAGAA-like motif sites reported by Kim *et al.* 2020 [43] shown in green; and **v)** microRNAs predicted from the observable sub-genomic RNAs in this study shown in brown. **B)** Non-random proximity of RBs and HSPs to ToCs in SARS-CoV-2. Violin plots showing nearest distances of the ToCs (n = 6) and other TFOs (n = 12) to actual RBs and HSPs and distances from the ToCs to randomly generated sites (random). Adjusted (FDR) *p-*values: *p* < .05 (*), *p* < .01 (**), *p* < .001 (***), Wilcoxon rank sum test. Error bars represent standard error of the mean. Genome-wide analysis shows breakpoints and hotspots are significantly closer to ToCs than other TFOs or random sites. **C)** ORF-constrained analysis confirms breakpoints and hotspots are significantly closer to ToCs than to other TFOs and from randomized sites.

### Enrichment of svRNA-TFO targets across infection-relevant transcriptomes

To identify potential targets of svRNA-mediated epigenetic interference during COVID-19, we identified 120 genes that were differentially expressed upon SARS-CoV-2 infection and overlapped with associated enhancer regions predicted to be targeted by the six svRNA-TFOs of concern (Supplemental Figure S6). This corresponds to a ∼ 2.5-fold enrichment over the background Differentially Expressed Gene (DEG) rate and is statistically significant (one-sided Fisher’s test, *p*-value < .001; Supplemental Figure S7A). Significant enrichment was also confirmed related to a TFO-based background simulated from 1,000 random enhancer-target sets (*p-*value < .001; Supplemental Figure S7B). Notably, 16 % of the DEG-matched enhancer gene targets (AHNAK, ALDH4A1, APOBEC3A, CD82, CHAC1, DLL4, EGFL7, FZD4, GAS6, IFNGR2, IL1R2, LILRB4, MYH10, NAPA, PLXNA2, SERPINB1, SPSB1, TNS1, and UGCG) were shared across two independent transcriptome studies included in our analysis, highlighting these as potentially the most plausible targets of svRNA-mediated epigenetic interference in lungs during infection of SARS-CoV-2 (Supplemental Figure S6, Supplemental Table S3).

### Analyses of six svRNA-TFOs in SARS-CoV-2 and SARS-CoV-1

Of the six putative svRNA-TFOs of concern identified in this study, the most conspicuous is svRNA-TFO-N, which partially overlaps with one of two dominant small RNA-seq high-coverage regions detected in both SARS-CoV-2 and SARS-CoV-1 (Figure 3A and B). In SARS-CoV-2, a marked reduction in small RNA-seq coverage is observed at nucleotides 29,376 – 29,492 compared to the region directly upstream (Figure 3A). The location of the abrupt drop in small RNA-seq coverage overlaps with two secondary modification sites, known as ‘AAGAA-like’ motif sites (Kim et al., 2020), which are located directly upstream and downstream of the TFO. The svRNA-TFO-N encodes a polypurine type TFO motif (5′-AAAAAGAAGAAGG-3′), which is predicted to have binding affinity for the enhancers of genes encoding the CREB protein (CREBBP), alanyl aminopeptidase (ANPEP), interferon stimulated exonuclease gene (ISG20), and fizzled class receptor 4 (FZD4). Expression of these genes was found to be significantly abrogated during infection of SARS-CoV-2 in COVID-19, human lungs, or Calu-3 cells based on previous transcriptome analyses (Supplemental Figure S6F).

**Figure 3:**
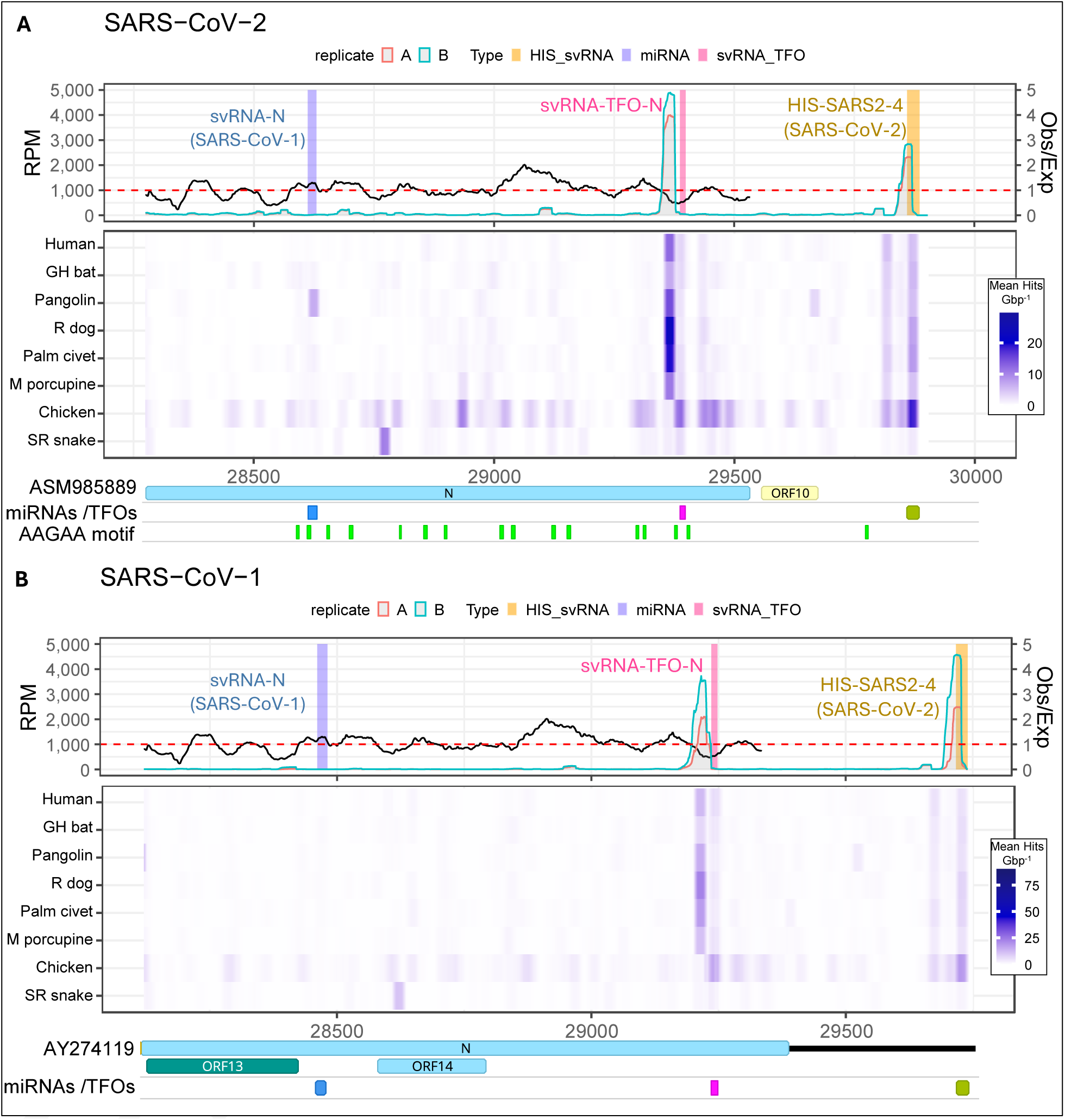
Nucleocapsid (N) + 3′ untranslated region integrating small RNA-sequencing (RNA-seq), synonymous-site conservation (SSC) analysis, microRNAs (miRNAs) and other annotations of interest, triplex-forming oligonucleotide (TFO) predictions, and host genome homology. Topmost plots, first *y*-axis: reads per million (RPM) coverage small RNA-seq from Calu-3 cells 12 h post-infection [45]. Second *y-*axis SSC analysis for *Sarbecovirus*, red dashed-line representing an equal ratio of observed (Obs) synonymous mutations compared to the number of expected (Exp). Middle heatmaps: rolling average BLAST hits against various animal genomes (GH bat = greater horseshoe bat; R dog = raccoon dog; M porcupine = Malaysian porcupine; and SR snake = San Diego ring-necked snake) (see Materials and Methods). Bottom features plots: TFOs predicted against human lung enhancer sequences and svRNA-TFOs of concern identified using a functional genomics pipeline in this study (Supplemental Table S1); HIS-SARS2-4 (SARS-CoV-2) was identified by Li et al. [7]; ‘AAGAA-like’ (AAGAA) motifs were identified by Kim et al. [43]; and svRNA-N (SARS-CoV-1) was identified by Morales et al. [4]. **A)** SARS-CoV-2 genetic region 28,274 – 29,903 in Wuhan-Hu-1 reference genome **B)** SARS-CoV-1 genetic region 28,120 – 29,750 in Tor2 reference genome.

Using Synplot2 to assess synonymous-site conservation (SSC) among *Sarbecovirus* genomes and recMRCA aligned sequences, we observed a pronounced increased in SSC across most of the svRNA-TFO-N region (Figure 3). Short 25-nt SARS-CoV-2 genome segments were also queried against animal genomes using BLASTn, revealing a discrete overlap between a mammalian-specific highly-homologous region (HHR) and the high-coverage small RNA-seq region corresponding to the same area (Figure 3A). Two additional regions in the 3′ UTR of the Wuhan-Hu-1 reference genome showed relatively high homology to animal genomes, including one HHR associated with the namiRNA HIS-SARS2-4, which partially overlaps the poly-A tail region [7]; these patterns were consistent in both SARS-CoV-2 and SARS-CoV-1 (Figure 3A and B). The TFO associated with svRNA-TFO-N is a conserved 12-nt polypurine motif (5′-AARAAGAARAAG-3′) (Supplemental Figure S8C) present in all *Sarbecovirus* genomes except RsYN04, which contains a cytosine at position 10 (Supplemental Figure S8A and B).

To explore effects of mutations associated with SARS-CoV-2 VoCs on putative svRNA-TFO progenitor structures, minimum free energy (MFE) of optimal and three-dimensional (3-D) RNA structure remodelling analyses were used. A 150 nt sequence was used to represent the svRNA-TFO-N progenitor from the Wuhan-Hu-1 strain, with a thermodynamically stable MFE of optimal structure containing multi-branched hairpins (Supplemental Figure S9A). The bases that had the most RPM in small RNA-seq coverage (29,343 – 29,376 in Wuhan-Hu-1) were found on an unpaired end within an area that was not predicted with high probability or reliability in either the MFE of optimal or 3-D structural models, respectively. However, in both models a higher-confidence hairpin spanning 29,422 – 29,463 was consistently observed (Supplemental Figure S9A and B). When comparing the predicted svRNA-TFO-N progenitor structure from Wuhan-Hu-1 against the Delta VoC (B1.617.2) that carries the U29402 (Y377) variant of the D377Y mutation, only a small increase in the Gibbs free energy (ΔG) of the MFE of optimal from -31.90 to -31.10 kcal mol^-1^ was found (Supplemental Figure S9A and C). Similarly, for the 3-D structural model, this mutation only had a minor effect on the overall reliability average, which slightly decreased from 65.8 to 64.2 (Supplemental Figure S9B and D). Using both the MFE of optimal and 3-D structural models, the G→U transition at position 29,402 is predicted to confer a change in the loop of a hairpin that is located downstream from the TFO, which is also a site for one the most common secondary modification known from SARS-CoV-2 [43] (Supplemental Figure S9).

Beyond the conspicuous svRNA-TFO-N, we also found interesting results in the nsp3 region among SARS-CoV-2 and SARS-CoV-1. When considering small RNA-seq coverage, SSC analysis and host-homology searches, it was revealed that SARS-CoV-2 and SARS-CoV-1 nsp3 regions both share similar polypurine TFO-like tracts but strikingly the small RNA-seq coverage and presence of an HHR within the vicinity of the TFO is clearly different between the two viruses (Figure 4A and B). The svRNA-TFO-nsp3 in the SARS-CoV-1 encodes a polypurine motif (5′-GAGGAAGAAGAAA-3′), which differs by 3 nt from the svRNA-TFO-nsp3 encoded by SARS-CoV-2 (5′-GAAGAAGAAGAG-3′) (see also Supplemental Figure S10). Using short BLASTn, a dominant HHR was found within the vicinity of a different TFO-like sequence in SARS-CoV-1, which overlaps with a region of high SSC (Figure 3B), as well as svRNA-nsp3.2 previously characterised by Morales et al. [4]. We did not find the same pattern in SARS-CoV-2 (Figure 3A), with the alignments in this region showing an insertion has occurred in the svRNA-nsp3.2 region among the SARS-CoV-2-related subgroup (Supplemental Figure S11). Notably, the svRNA-TFO-nsp3 in SARS-CoV-2 is predicted to interact with only three lung enhancer gene targets, *HADH*, *TMED2*, and *COL16A1*, which are also differentially expressed in Calu-3 cells [45]. Comparatively, this is a much lower number of lung enhancer gene targets than the other putative svRNA-TFOs identified in this study, which ranged from 6 – 19 (Supplemental Figure S6B).

**Figure 4:**
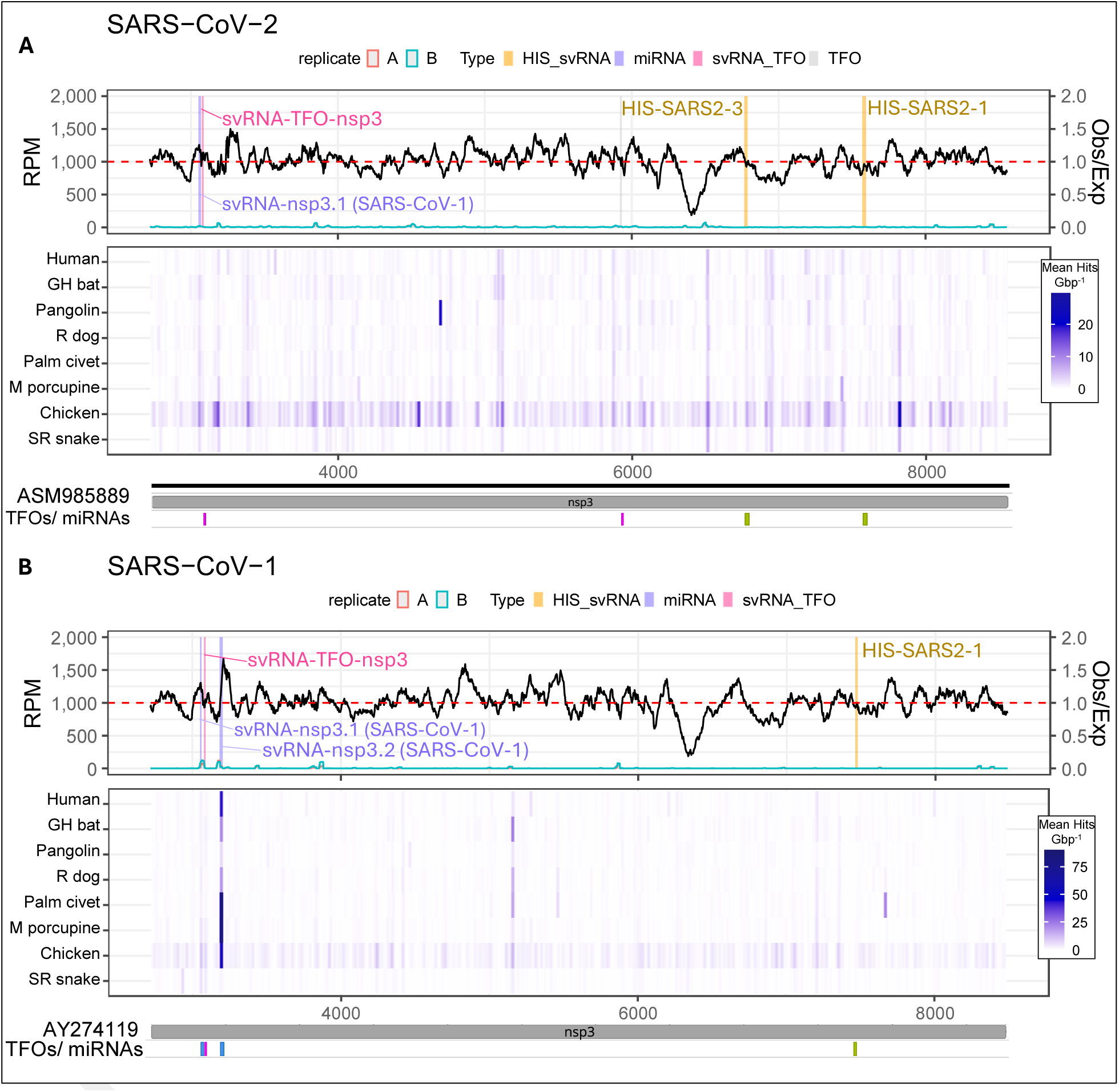
nsp3 region integrating small RNA-sequencing (RNA-seq), synonymous-site conservation (SSC) analysis, microRNAs (miRNAs) and other annotations of interest, triplex-forming oligonucleotide (TFO) predictions, and host genome homology. Topmost plots, first *y*-axis: reads per million (RPM) coverage small RNA-seq from Calu-3 cells 12 h post-infection [45]. Second *y*-axis: SSC analysis for *Sarbecovirus*, red dashed-line representing an equal ratio of observed (Obs) synonymous mutations compared to the number of expected (Exp). Middle heatmaps: rolling average BLAST hits against various animal genomes (GH bat = greater horseshoe bat; R dog = raccoon dog; M porcupine = Malaysian porcupine; and SR snake = San Diego ring-necked snake) (see Materials and Methods). Bottom feature plots: TFOs predicted against human lung enhancer sequences and svRNA-TFOs of concern identified using a functional genomics pipeline in this study (Supplemental Table S1); HIS-SARS2-1 and HIS-SARS2-3 were identified by Li et al. [7]; svRNA-nsp3.1 (SARS-CoV-1) and svRNA-nsp3.2 (SARS-CoV-1) were identified by Morales et al. [4]. **A)** SARS-CoV-2 genetic region 2,720 – 8,554 in Wuhan-Hu-1 reference genome. **B)** SARS-CoV-1 genetic region 2,719 – 8,484 in Tor2 reference genome.

Also, two potential triplex-forming svRNAs were identified from the S-ORF regions using the Epi-VIRTEX approach (Supplemental Figure S12). svRNA-TFO-S.1 and svRNA-TFO-S.2 were identified as being weakly expressed in Calu-3 cells by SARS-CoV-2 at 24 hpi (Figure 1C-D) compared to svRNA-TFO-N (Figure 1F) and svRNA-TFO-nsp2 (Figure 1A) based on the relative levels of small RNA-seq coverage. The TFOs of svRNA-TFO-S.1 and svRNA-TFO-S.2 are in regions of slightly lower-than-expected SSC and the TFO of svRNA-TFO-S.1 shares the same location with an HHR specific to some mammals (not Malaysian porcupine) in SARS-CoV-2, but not in SARS-CoV-1 (Supplemental Figure S12). The svRNA-TFO-S.1 TFO sequence is conserved among *Sarbecovirus*, consisting of a mixed purine-pyrimidine (5′-UUUUGGUGGUGU-3′) motif (Supplemental Figure S13A-C). Also, the TFO sequence of svRNA-TFO-S.2 a conserved mixed purine-pyrimidine type motif, but its motif contains a palindrome (5′-UUUUGGUGGUUUU-3′) (Supplemental Figure S14A-C).

Putative human lung enhancer gene targets for svRNA-TFO-S.1 include genes for two antiviral C→U RNA-editing cytidine deaminases APOBEC3A and APOBEC3F, and a non-muscle myosin heavy chain subcomponent II-B (MYH10) that could be important during infection (Supplemental Figure S6C). The enhancer gene targets of svRNA-TFO-S.2 include several with immune-related functions such as, interleukin 1 receptor type 2 (IL1R2), serpin family B member 1 (SERPINB1), and leukocyte immunoglobulin-like receptor subfamily B member 4 (LILRB4) (Supplemental Figure S6D).

Using RNA structural remodelling of the S:D614G mutation in the putative svRNA-TFO-S.1 progenitor RNA structure, the G23403 variant (characteristic of Alpha [B.1.1.7], Beta [B.1.351], Gamma [P.1], Delta [B.1.617.2], and Omicron [B.1.1.529]) results in a side bulge expansion plus a substantive decrease in the structure’s MFE of optimal (-30.19 kcal mol^-1^) when comparing the same structure predicted for Wuhan-Hu-1 (-22.51 kcal mol^-1^) (Supplemental Figure S15 A and C). The increased stability of the G23403 variant is likely due to the avoidance of an unpaired adenine adjacent to the side bulge. Noting that 3-D modelling of the putative svRNA-TFO-S.1 progenitor resulted in an overall lower reliability model compared to that of the svRNA-TFO-N (Supplemental Figure S9B and D); however, the G23403 variant was predicted to exert effects on the orientation of a side-bulge similar to the MFE of optimal model for svRNA-TFO-S.1 progenitor structure, as well as having effects on the lower-reliability 5′ and 3′ ends (Supplemental Figure S15B and D).

While an overlapping HHR to the TFO of svRNA-TFO-E in SARS-CoV-2 and SARS-CoV-1 was detected, this HHR is not mammalian-specific, and the region shared limited SSC among *Sarbecovirus* compared to the other svRNA-TFOs mentioned above (Supplemental Figure S16A,B). The TFO associated with svRNA-TFO-E consists of a 12-nt long poly-pyrimidine motif (5′-CUUCUUUUUCUU-3′) that is conserved among *Sarbecovirus* (Supplemental Figure S17A,B,C). Potential enhancer-gene targets of svRNA-TFO-E include genes for host cytokine signalling and immune response to viruses, such as interleukin 1 receptor type 2 (IL1R2), interferon gamma receptor 2 (IFNGR2), and tetraspanin CD82 (Supplemental Figure S6E). Also, the gene for SNARE-associated NSF attachment protein alpha (NAPA), which is involved in membrane-vesicle fusion, is another potential enhancer-associated target gene of svRNA-TFO-E.

Based on the small RNA-seq coverage in the nsp2 region, SARS-CoV-2 has several areas of > 150 RPM coverage compared to SARS-CoV-1, including one high-coverage area adjacent to the TFO from this region (Supplemental Figure S18A, B), noting also that the polypurine motif (5′-AAAAAGAGAAAG-3′) is exclusive to the SARS-CoV-2-related sub-group (Supplemental Figure S19A,B,C). Our analysis identified potential enhancer-gene targets for the svRNA-TFO-nsp2 against human lungs, including genes for a growth-arrest specific protein (GAS6), CREBBP, and neuroblast differentiation-associated protein (AHNAK) (Supplemental Figure S6A). Lastly, the nsp2 region was not found to contain any major mammal-specific HHRs, and there was limited SSC associated with high-coverage/TFO region (Supplemental Figure S18).

## DISCUSSION

By integrating RNA-seq data, evolutionary models, and sequence analysis, we propose a potential role for svRNA-mediated epigenetic interference in the evolution of SARS-CoV-2 and SARS-CoV-1 within the *Sarbecovirus* lineage. Building on prior findings that SARS-CoV-2 abundant svRNAs can be detected in small RNA-seq [3] that can serve as early biomarkers of infection [67], our analysis of direct and small RNA-seq data from SARS-CoV-2-infected cells [43], combined with TFO detection, identified six putative svRNA-TFOs of concern. Notably, two of these, svRNA-TFO-N and svRNA-TFO-nsp2, are produced in high abundance as early as 12 hpi in Calu-3 cells [45]. We also find highly significant enrichment of the putative svRNA-TFO enhancer-associated gene targets from infection-relevant host DEGs, which supports the epigenetic interference as viable mechanism in SARS-CoV-2. Furthermore, predictive structural modelling of the svRNA progenitors for svRNA-TFO-N and svRNA-TFO-S.1 suggests a potential link to respective mutations that became characteristic in later SARS-CoV-2 VoCs, including S:D614G, and N:D377Y.

Importantly, all six of the putative svRNA-TFOs of concern identified in this study were significantly co-located within 183 nt of approximately 26 % of recombination breakpoints hotspots in the SARS-CoV-2 genome [66]. Comparable patterns have also previously been observed in other viruses. In herpes simplex virus 1 (HSV-1), guanine-rich G-quadruplex structures lie within 500 nt of 11 % of recombination breakpoints [68,69], and in human immunodeficiency virus (HIV), envelope-gene breakpoints consistently co-localize with structured RNA regions and areas of high sequence conservation [70]. Together, these parallels point toward a broader principle in viral genome biology, namely that conserved RNA structures and sequence-constrained regions can influence recombination landscape. Consistent with this, several of the svRNA-TFO regions of concern in our study displayed significant synonymous-site conservation across the *Sarbecovirus*, suggesting a deeper evolutionary role for svRNA-TFOs in shaping genome architecture within this lineage. Future molecular studies are required to validate these findings.

Homology between regions of the SARS-CoV-2 and human genomes has been described before also [7,67], including with the functional characterization of five human identical sequence (HIS) namiRNAs produced by SARS-CoV-2 that bind to enhancer sequences and control host gene expression [7]. Our study goes a step further by expanding virus-to-host homology searches against the reference genomes of various mammalian hosts and non-mammalian controls. Strikingly we show discrete regions of higher-than-average homology specific to mammalian host genomes for svRNA-TFO-N and svRNA-TFO-S.1. While the latter was specific to SARS-CoV-2, another discrete high homology region with a less clear host-matching pattern was identified in the nsp3 ORF of SARS-CoV-1. Together these results broadly suggest a potential role for epigenetic interference in host-adaptation in *Sarbecovirus*, which requires further attention.

In another SARS-CoV-2 sequencing study, researchers used *in situ* conformation sequencing to map viral genome structure [71]. These results aligned with our model that the A23403G (S:D614G) mutation stabilizes the svRNA-TFO-S.1 progenitor RNA structure by creating a more thermodynamically favourable six-nucleotide side bulge 56-nt downstream of the TFO of concern, which was also consistent with strong SHAPE-MaP signals from infected cells [72]. Beyond its predicted RNA effects, the G614 spike variant is also known to increase infectivity [73], transmissibility [74], and virion stability [75], supporting the ideas that G23403 provided dual advantages, improving both spike protein function and svRNA-TFO-S.1 progenitor stability, at a pivotal point early in SARS-CoV-2 evolution. This is consistent with G23403 rising to fixation during an eight-month period of evolutionary stasis beginning in April 2020 and persisting across all subsequent lineages that began diverging in August 2021 [76]. A second mutation in the nucleocapsid region, G29402U (N:D377Y), associated with the Delta variant also showed RNA-structural effects in our modelling, altering a stem loop in the MFE of optimal structure and producing subtle effects in the corresponding 3-D model. G29402U co-occurs with G28881U (N:R203M), and together these mutation enhance viral suppression of RIG-I-mediated antiviral signalling [77]. The G→U transition at 28,881 additionally introduces a premature AUG (Met) codon into the nucleocapsid sgRNA located ∼500 nt upstream from the start of the svRNA-TFO-N region. This suggests that both G29402U and G28881U may influence svRNA-TFO-N processing, although this requires further confirmation with small RNA-seq comparing svRNA-TFO-N production levels between the Delta variant and Wuhan-Hu-1.

Notably, A23403G (S:D614G) is consistently accompanied by three further mutations (C241U, C3037U, and C14408U) [71], all of which are C→U transition; intriguingly, svRNA-TFO-S.1 is predicted to target enhancer regions of *APOBEC3A* and *APOBEC3F*, which are genes for cytidine deaminases that drive C→U conversions and contributes to the known C→U mutational bias in SARS-CoV-2 [78,79]. This raises the possibility that svRNA-TFO-S.1 could have been positively selected if APOBEC upregulation increased the emergence of novel RNAse L cut-sites, an idea consistent with the central role of RNAse L in SARS-CoV-2 [53–56] and its preference for unpaired UU or UA dinucleotide in stem-loops bulges [80,81]. Supporting this model, Cao et al. [71] found that all three accompanying mutations (C241U, C3037U, and C14408U) introduce new unpaired UU dinucleotides specifically into bulge regions across three distinct stem-loop structures, offering a mechanistic link between APOBEC-driven editing, svRNA-TFO function, and structural evolution of the SARS-CoV-2 genome.

Another major highlight of this study is that, beyond the six putative svRNA-TFOs of concern identified in SARS-CoV-2, two previously characterized svRNAs, a canonical miRNA (svRNA_nsp3.2) from SARS-CoV-1 [4] and a namiRNA (HIS-SARS2-4) from SARS-CoV-2 [7], also appear plausibly linked to epigenetic interference. The nsp3 region in both SARS-CoV-1 and SARS-CoV-2 contains multiple TFO-like polypurine motifs that cluster near a known recombination breakpoint [66], strongly suggesting this locus has an evolutionary history involving recombination across the *Sarbecovirus* lineage. Our analysis indicates that svRNA-nsp3.2 may participate in epigenetic interference, as it overlaps one of these TFO-like motifs, is produced at high abundance in Calu-3 cells and has a HHR against several mammalian host genomes unique to SARS-CoV-1. Beyond the nsp3 region, converging evidence from small RNA-seq coverage, SCC analysis, and cross-species genome homology indicates the production of two highly abundant putative svRNAs derived from the N-ORF and 3′ UTR in *Sarbecovirus*, including svRNA-TFO-N exhibiting the single highest small RNA-seq coverage in the SARS-CoV-2 genome across multiple studies [3,67] and the HIS-SARS2-4, a namiRNA encoded in the 3′ UTR that controls expression of *hyaluronan synthase 2* during COVID-19 [7]. The close proximity of HIS-SARS2-4 and svRNA-TFO-N (within ∼ 550 nt of each other) suggests a shared processing mechanism for nuclear-acting svRNAs and svRNA-TFOs. This interpretation is further supported by the exceptionally high transcriptional activity of this particular genomic region, driven by discontiguous transcription and the disproportionately high production of the nucleocapsid sgRNA, relative to other sgRNAs from SARS-CoV-2 [43,45]. Together, these observations point to a broader network of svRNAs, including svRNA-TFO-N, svRNA-nsp3.2, HIS-SARS2-4, and the five additional svRNA-TFOs identified here. These svRNAs may act synergistically in epigenetic interference, an area that warrants dedicated experimental investigation.

Lastly, the application of advanced RNA-sequencing technologies has created unprecedented opportunities to interrogate virus evolution. When integrated with evolutionary analyses and RNA structural modelling, as demonstrated in this study, these datasets can uncover previously unrecognized mechanisms, including svRNA-mediated epigenetic interference in SARS-CoV-2. Although the present study was deliberately focused on datasets relevant to SARS-CoV-2 infection of human lungs and lung-derived cell types and tissue as a biologically grounded starting point, extending these analyses to additional infection-relevant cell types and tissues, together with direct molecular evidence, will be essential to define the broader aspects of this mechanism. Such efforts are likely to refine our understanding of the natural drivers of *Sarbecovirus* evolution and inspire innovative antiviral strategies. Ultimately, these advances may enhance preparedness for potential future outbreaks of SARS-CoV-like viruses, which continue to pose major threats to human health and global stability.

## MATERIALS AND METHODS

### Triplex-forming oligonucleotide and microRNA prediction in SARS-CoV-2

To investigate epigenetic interference as a potential virulence mechanism in SARS-CoV-2, putative triplex-forming oligonucleotides (TFOs) from the Wuhan-Hu-1 reference genome were identified from the human lung-associated enhancer sequences from EnhancerAtlas 1.0 [59] (downloaded May 29, 2020) using *Triplexator* [58] (Virtual Box v4.2.12; build 84980, downloaded January 23, 2021). The *Triplexator* algorithm implements sequence-matching based on canonical triplex-formation rules and was implemented using the lowest allowable size limit of 12 nt.

Next, sequences representing 477 observed SARS-CoV-2 sgRNAs during infection of Vero Cells [43] were obtained from the University of California, Santa Cruz (UCSC) SARS-CoV-2 Genome Browser [82,83] (downloaded on September 8, 2025 [http://genome.ucsc.edu/cgi-bin/hgTables]). By collapsing redundant sequences at a 99 % identity threshold using CD-HIT (v4.8.1-2019-0228), 153 non-redundant sgRNA sequences were carried forward for further analysis [84]. Using SeqKit (v0.12.1) [85], these sequences were fragmented with a window-size and sliding-step length of 120 nt and 20 nt, respectively. Each fragmented sequence was then analysed for its structural thermal stability with ViennaRNA (v2.4.14) *RNAfold* [86] to identify 5,670 structures meeting the -20 kCal mol^-1^ cut-off threshold for predicted minimum free energy (MFE) of optimal. From these RNA structures, 223 pre-miRNA hairpins were predicted by the machine learning tool HuntMi (downloaded May 23, 2020) trained on a virus-specific geometric mean-optimised training set [87]. Lastly to predict mature miRNAs, a naïve Bayes classifier MatureBayes was used to identify 446 mature miRNAs from SARS-CoV-2 [88] (http://mirna.imbb.forth.gr/MatureBayes.html; accessed September 22, 2025). Non-redundant representatives of these miRNAs, along with 331 other previously predicted SARS-CoV-2 miRNAs from genome-based studies [61–65] were mapped to the Wuhan-Hu-1 reference genome to identify putative svRNA-TFOs of concern, along with sufficient small RNA-seq coverage (see below). A number of miRNAs or nuclear-activating miRNAs (namiRNAs) functionally characterised in SARS-CoV-2 [5–8] and SARS-CoV-1 [4] were also mapped against the respective reference genomes (Wuhan-Hu-1 or Tor2) and used for comparative purposes with small RNA-seq.

### Small RNA-sequencing coverage and normalization

To identify potential svRNA-TFOs being produced within 24 hpi, a small RNA-seq a cell-based experiment using Calu-3 lung cancer epithelial cells was combined with the miRNA predictions from this, and other studies, for the SARS-CoV-2 Wuhan-Hu-1 genome described above. The small RNA-seq data collected at 4, 12, and 24 hpi were downloaded from NCBI’s GeoBank (GSE148729) on November 15, 2021 (SARS-CoV-2) and January 21, 2024 (SARS-CoV), along with the corresponding mock-infected and uninfected controls [45]. Following a similar two-pass trimming approach described in in the originating paper but instead using Flexbar (v3.0.3) [89], the first, 3 nt on the 5′ end were removed, the 3′ end was trimmed to remove bases with a quality score of < 30, and the Illumina small TruSeq adapter was removed from the 3′ end. On the second-pass, the polyA adapter was removed if there was an overlap of 10 nt with no mis-matches. Trimmed/filtered sequences were next aligned to the SARS-CoV-2 reference genome (NC_045512.2; Wuhan-Hu-1) using Bowtie2 (v2.3.4.1) [90].

Alignment files were sorted using Samtools (v1.10) [91] and RPM coverage was determined with bedtools (v2.27.1) [92]. For visualization purposes, ggplot2 (v3.3.3) [93] was used to show coverage of two biologically-replicated libraries over the presumed svRNA-TFO progenitor RNA structure regions (see methods below), or else the coverage plots were centred on the TFO. To identify the potential svRNA-TFOs of concern, regions encoding predicted TFOs were assessed for small RNA-seq coverage. While previous small RNA-seq studies detecting miRNAs have used RPM thresholds ranging from 10 to 100 [94–96], we applied a conservative cut-off for identifying SARS-CoV-2 svRNAs. Candidates were required to reach ≥ 40 RPM within 25 nt of the predicted TFOs in both experimental replicates at 12 or 24 hpi. For comparing the level of RPM small RNA-seq coverage between SARS-CoV-1 and SARS-CoV-2, analysis focused on the 12 hpi timepoint because of the reported difference in the number of viral particles between the SARS-CoV-2- and SARS-CoV-infected Calu-3 cells at 24 hpi, and that the coverage was relatively lower at 4 hpi as expected [45].

### Phylogenetic reconstruction, synonymous-site conservation analysis, motif searches, and recombination breakpoint and hotspot analysis

Using the R package *DECIPHER* (v2.26.0) [97], amino acid-based multiple sequence alignments (MSAs) were made from 76, 85, 80, 54, and 76 non-redundant translated nucleotide sequences derived from the nsp2, nsp3, S, E, and N ORF regions, respectively. The sequences used in this analysis included SARS-CoV-2 (Wuhan-Hu-1) and SARS-CoV-1 (Tor2) reference genomes, and the genomes of 96 other bat- and pangolin-infecting *Sarbecovirus* available from NCBI’s GenBank and the GISIAD repositories (Supplemental Table S4). This analysis also included sequences derived from SARS-CoV-2’s most-recent reconstructed ancestor (MRCA) [98]. Sequences containing ambiguous bases were not included in the alignments.

To improve alignment of the nsp3 ORF region, the MRCA nsp3-associated sequence was aligned secondarily using Clustal Omega (v1.2.2) with default settings in Geneious Prime® (2023.2.1). To achieve the S-ORF alignment, sequences derived from the following accessions were omitted: KF294457.1, DQ412043.1, DQ648857.1, KJ473815.1, and NC_014470.1. Following multiple sequence alignment, the nsp2, nsp3 and S-ORF regions were partitioned according to the location of previously identified recombination breakpoints in Wuhuan-Hu-1 [66]. Next using *IQ-TREE 2* (v1.61) [99], phylogenetic analysis was performed on the aligned E-and N-ORF and partitioned nsp2, nsp3, and S-ORF sequences by best-fit substitution model selection according to the Bayesian Information Criterion. Node confidence was tested using 10,000 ultrafast bootstraps for each of the phylogenies generated.

Next the tip orders were extracted from each of the phylogenies to implement *Synplot2* (v2014) analysis for the nsp2, nsp3, S-, E-, and N-ORF regions, along with the amino acid-based transcribed alignments. *Synplot2* is an algorithm designed to analyse protein-coding regions from RNA viruses to identify statistically significant reductions in the variability at synonymous sites as an indicator for overlapping functional non-coding elements [100]. Also, segments of the alignments containing TFOs from the six svRNA-TFOs of concern were analyzed using the Multiple Em for Motif Elicitation (MEME) Suite (v5.5.7, v5.5.9) [101]. To identify conserved motifs with MEME, either all *Sarbecovirus*-related sequences were analyzed together (svRNA-TFO-N, svRNA-TFO-S.1, svRNA-TFO-S.2, svRNA-TFO-E, svRNA-TFO-nsp3) or separately among SARS-CoV-2-related (svRNA-TFO-nsp2). The phylogenies were rooted with using BtKY72 (KY352407.1), except the E-ORF was rooted with RmYN05 (MZ081376.1) for visualization purposes with the TFO-like sequence alignments using ggtree (v3.6.2) [102] and ggmsa (v1.4.0) [103] R packages, respectively.

Genome-wide and ORF-constrained permutation testing with pairwise Wilcoxon rank sum test and FDR multiple test correction [104] was used to test for a non-random association between the six putative svRNA-TFOs identified in this study and the previously identified recombination breakpoints (n = 21) and hotspots of 475 nt length (n = 2) in the Wuhan-Hu-1 genome (NC_045512.2) [66]. Base R (version 4.2.1) was used to randomly generate 1,200 sites, or the number of sites equal to 10 % of the sequence length for the genome-wide and ORF-constrained analysis, respectively. For the ORF-constrained analysis, nearest distance measures were normalised per kb length for each respective region.

### Short BLASTn-based host genome homology search strategy

High homology regions (HHRs) were identified in the genomes of SARS-CoV-1 and SARS-CoV-2 by comparing them to the human reference genome (GRCh38) and genomes of other mammals known to carry *Sarbecovirus*. This includes the Chinese pangolin (*Manis pentadactyla*), masked palm civet (*Paguma larvata*), and greater horseshoe bat (*Rhinolophus ferrumequinum*) (Supplemental Table S5). We also examined the genomes of the Malaysian porcupine (*Hystrix brachyura*) and common raccoon dog (*Nyctereutes procyonoides*), which are potential host species linked to SARS-CoV-2-postive PCR samples from the Huanan market in Wuhan, China [105,106]. The chicken (*Gallus gallus*) and San Diago ring-necked snake (*Diadophis punctatus similis*) served as control hosts.

To prepare for BLASTn-based short searching (v2.6.0+) [107], we segmented the reference genomes of SARS-CoV-2 (Wuhan-Hu-1; NC_045512.2) and SARS-CoV-1 (Tor2; AY274119.3) using SeqKit (v0.12.1) [85], with a window length of 25 nt and a single-nucleotide step. We identified hits with 100 % identity, an e-value threshold of 0.8, a minimum word-size of 7, a gap-open penalty of 5, and a gap-extend penalty of 2. We normalized the total hits by the total length in gigabase pairs (Gbp) for each animal host genome and visualized the results as a heatmap using ggplot2 (v3.4.4) [93] with a rolling average window size of 20 nt.

### Human enhancer target gene prediction and enrichment analysis

To identify human enhancer target genes that would be most likely affected by the six putative svRNA-TFOs of concern identified in this study, three SARS-CoV-2-relevant DEG sets were used, including the top 500 most up- and down-regulated genes in SARS-CoV-2-infected Calu-3 cells and human lungs [45,108] (noting Wyler et al. 2021 dataset was also analyzed for small RNA-seq coverage described above), and the 1,730 most upregulated genes from whole blood transcriptome of patients with severe COVID-19 compared to healthy donors [109]. The DEG lists from these studies (GSE148729, GSE147507, and GSE171110) were accessed through the “COVID-19-related gene sets (2021)” using *enrichR* (https://maayanlab.cloud/ covid19/collection/7; accessed July 4, 2024, and October 4, 2025) [110,111].

Enhancer gene targets for all 18 TFOs identified in this study were converted from Ensembl gene IDs (ENSGs) to UniProtKB universal gene identifiers using the UniProt Retrieve/ID mapping tool (uniprot.org; accessed April 14, 2023, and October 4, 2025) [112]. Enrichment analyses focusing on the DEGs and enhancer gene targets with matching gene symbols were then performed in Base R (version 4.2.1) using both non-parametric and parametric statistical methods. These included including a one-sided Fisher’s exact test comparing 120 of 350 observed enhancer gene targets associated with the six putative svRNA-TFOs to the expected background detection rate of 13.5 %, derived from combined transcriptome studies (4,287 out of 31,698 transcripts). In addition, observed enrichment was compared against a simulated TFO background by generating 1,000 random enhancer gene target sets of equivalent size from the 12 remaining TFOs identified in this study, with significance assessed using a standard normal distribution test (*Z*-score probability).

### Small viral RNA progenitor structural predictions and mutation remodelling

To determine the mostly likely svRNA-TFO progenitor RNA structure sequences, consensus among locations for 106 conserved RNA structured regions [113], the 777 miRNA predictions from SARS-CoV-2 Wuhan-Hu-1 (described above), and the regions of small RNA-seq coverage were sought as required. Resultingly, the putative progenitor structure of the svRNA-TFO-N was expanded to 150-nt length to accommodate for the extremely high-coverage region upstream of the TFO. Due to the heuristic nature of svRNA-TFO progenitor sequence selection, the resultant RNA structural predictions are recommended as preliminary models only, which require further laboratory testing to confirm the exact sequence length or structural elements yet to be determined.

To evaluate potential effects of VoC-associated mutations on svRNA-TFO-related RNA structures, the putative progenitor structures corresponding to svRNA-TFO-N and svRNA-TFO-S.1 were remodelled for the G29402U (N:D377Y) and A23403G (S:D614G) mutations, respectively. MFE of optimal represented by Gibbs free energy (ΔG) was used to predict the secondary structure of 120-nt long svRNA-TFO-S.1 progenitor (from 23,304 – 23,423 in Wuhan-Hu-1) at 37 °C using the Andronescu model [114] assuming GU pairs and circular molecule, plus with an additional energy contribution for coaxial stacking of helices using *RNAfold* on ViennaRNA webserver (v2.6.3) (accessed December 26, 2025) [86]. A similar procedure, but with the Turner model [115] without assuming circular molecule was used to predict the MFE of optimal structure for the svRNA-TFO-N progenitor (from 29,343 – 29,492 in Wuhan-Hu-1) (accessed December 30, 2024). The MFE of optimal-based structures were visualized using Geneious Prime® (v2023.2.1) with the partition function for pairing probabilities and the same parameters described above, and the overall MFE of optimal was calculated.

Next, 3-D models were generated for the putative svRNA-TFO-N and svRNA-TFO-S.1 progenitors using the trRosettaRNA webserver (accessed December 30, 2024, and December 16, 2025) [116] with the *RNAfold*-generated MFE of optimal structure as custom secondary structures. The resultant 3-D models were visualized in FirstGlance Jmol (V4.31) [117] (accessed February 2, 2025, and December 26, 2025), including base reliability estimates assigned using the Local Distance Difference Test according to the trRosettaRNA webserver: very high ≥ 90; high 70-90; medium 50-70; and low < 50.

### Data availability statement

This study did not generate any new sequencing data. All data needed to evaluate the conclusions in the study are reported in the Materials and Methods and supplementary materials, with the exception of viral sequences sourced from GISEAD for which the accessions are provided in Supplemental Table S4. See https://github.com/damselflywingz/EpiVIRTEX for open-access to the scripts used in this analysis.

## Author contributions

**Amber R. Paulson**: Conceptualization; data curation; formal analysis; investigation; methodology; visualization; writing – original draft; funding acquisition **Vincent Montoya**: Conceptualization; methodology; software; writing – review & editing; **Jeffrey B. Joy**: Conceptualization; methodology; resources; supervision; writing – review & editing.

