## Supplemental Information for "Functional genomic analysis reveals mechanisms of epigenetic interference in SARS-CoV-1 and SARS-CoV-2"

*List of Tables and Figures*

**Supplemental Table S1.** Detection of triplex-forming oligonucleotides (TFOs) of concern in SARS-CoV-2 using microRNA (miRNA) predictions and small RNA-sequencing (RNA-seq)*.

| TFO position | # Triplex target sites | TFO type | Open reading frame | Overlap with any predicted miRNA precursor (see supplemental Table S2)? | ≥ 40 RPM Small RNA-seq coverage within 25 nt of TFO? |
|---|---|---|---|---|---|
| 1,130 – 1,141 | 79 | GA | nsp2 | Kahn et al. 2020 | No |
| 1,629 – 1,640 | 48 | GA | nsp2 | This study (adjacent) | Yes |
| 2,072 – 2,083 | 91 | GU | nsp2 | Kahn et al. 2020; Liu et al. 2021 | No |
| 3,074 – 3,085 | 7 | GA | nsp3 | Saini et al. 2023 | Yes |
| 5,920 – 5,932 | 133 | GU | nsp3 | | No |
| 9,247 – 9,258 | 106 | GU | nsp4 | This study; Kahn et al. 2020 | No |
| 11,174 – 11,187 | 527 | GU | nsp6 | This study | No |
| 11,795 – 11,810 | 635 | GU | nsp6 | Liu et al. 2021 | No |
| 15,726 – 15,740 | 80 | GU | nsp12 | Kahn et al. 2020 | No |
| 19,916 – 19,927 | 110 | GU | nsp15 | | No |
| 21,731 – 21,742 | 38 | CU | S | This study | No |
| 21,980 – 21,992 | 97 | GU | S | Kahn et al. 2020 | No |
| 23,335 – 23,346 | 81 | GU | S | This study | Yes |
| 23,950 – 23,962 | 160 | GU | S | This study | Yes |
| 25,643 – 25,655 | 102 | GU | ORF3a | | No |
| 25,700 – 25,711 | 130 | CU | ORF3a | | No |
| 26,296 – 26,307 | 53 | CU | E | This study; Kahn et al. 2020 | Yes |
| 29,387 – 29,399 | 101 | GA | N | This study | Yes |

*Small RNA-seq data from SARS-CoV-2 during infection of Calu-3 cells measured in reads per million (RPM), see Wyler *et al.* 2021 [1].

**Supplemental Table S2.** Predicted microRNA annotations for SARS-CoV-2 Wuhan-Hu-1 used in this study. See Materials and Methods for more information about the referenced studies.

20252531_Supplemental_Table_S2.xlsx

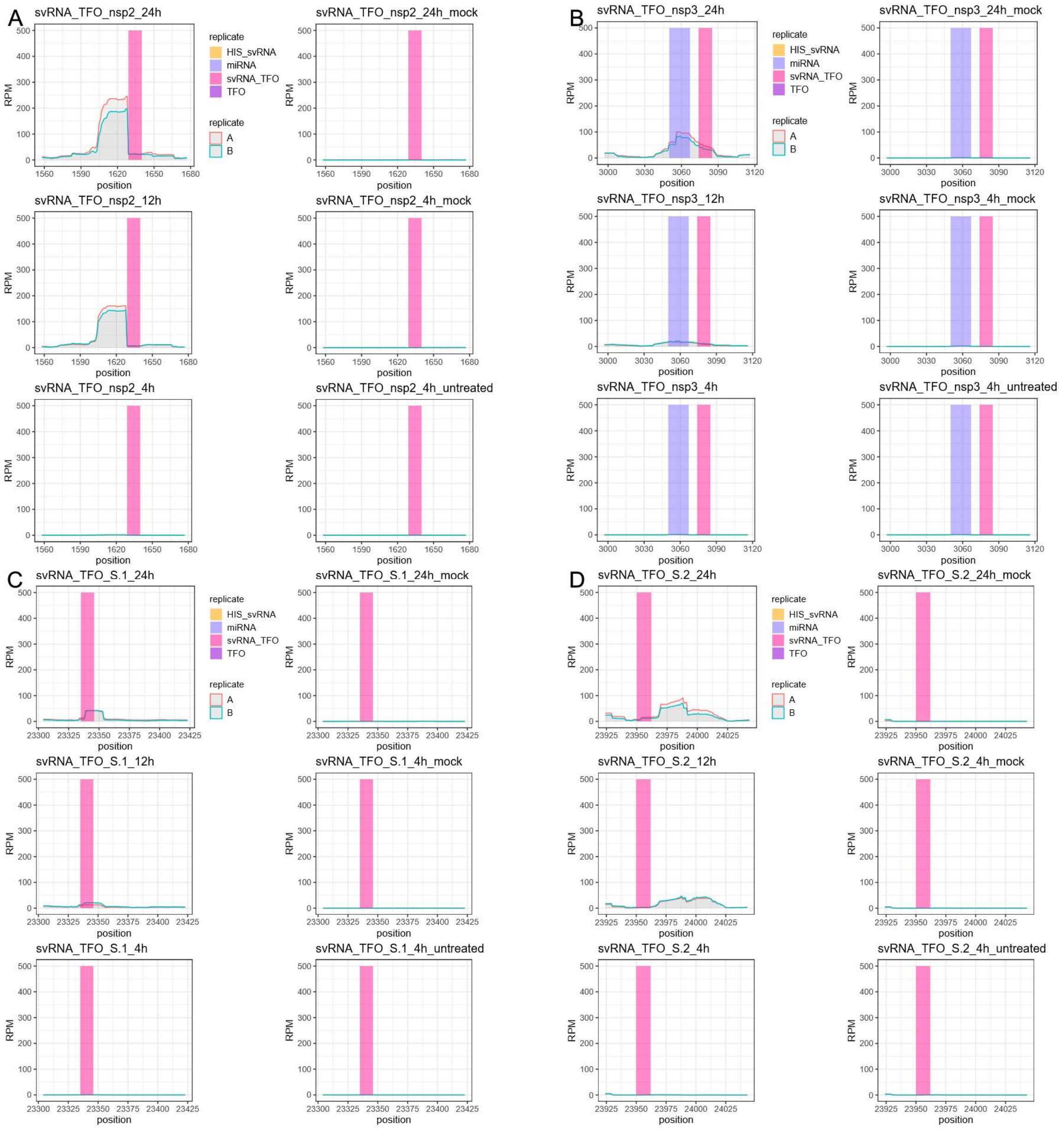

**Supplemental Figure S1. Single-nucleotide coverage of small RNA-sequencing reads (A and B replicate) from SARS-CoV-2-infected Calu-3 cells at 24, 12, and 4 hpi, with mock-infected and untreated as controls** [1]. **A)** portion of the nsp2 ORF region (1,558 – 1,677) encoding predicted svRNA-TFO-nsp2. **B)** portion of nsp3 ORF region (2,997 –3,116) encoding predicted svRNA-TFO-nsp3. **C)** portion of spike ORF region (23,304 – 23,423) encoding predicted svRNA-TFO-S.1. **D)** portion of spike ORF region (23,924 – 24,043) encoding predicted svRNA-TFO-S.2. The y-axis is presented as the number of reads per million (RPM) and all positions are relative to the Wuhan-Hu-1 genome.

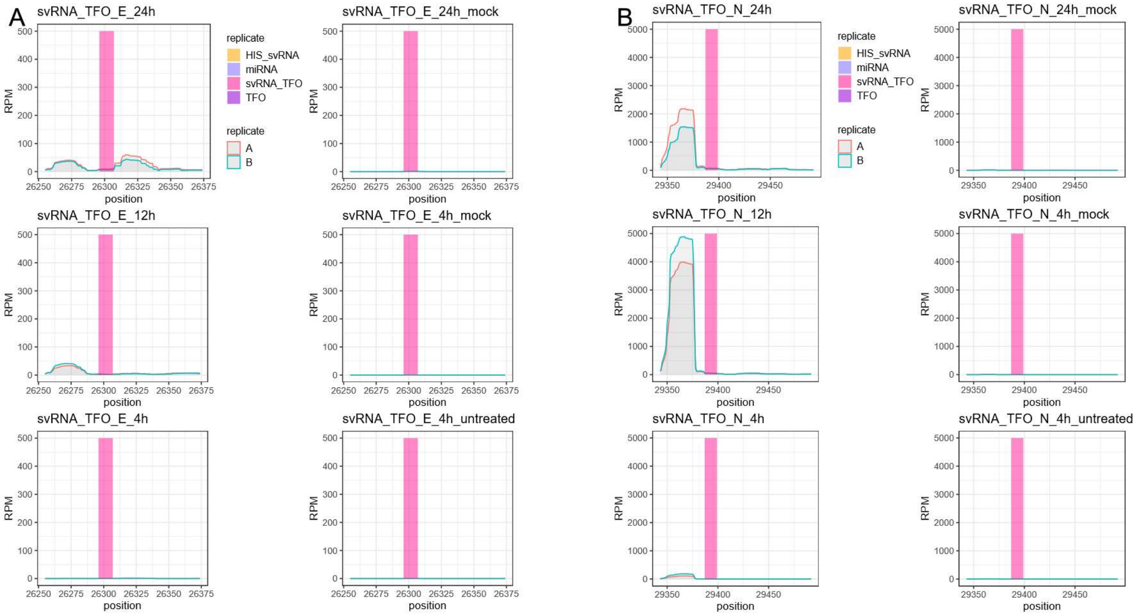

**Supplemental Figure S2. Single-nucleotide coverage of small RNA-sequencing reads (A and B replicate) from SARS-CoV-2-infected Calu-3 cells at 24, 12, and 4 hpi, with mock-infected and untreated as controls** [1]. **A)** portion of the envelope ORF region (26,255 – 26,374) encoding predicted svRNA-TFO-E. **B)** portion of the nucleocapsid ORF region (29,343 – 29,492) encoding predicted svRNA-TFO-N. The y-axis is presented as the number of reads per million (RPM) and all positions are relative to the Wuhan-Hu-1 genome.

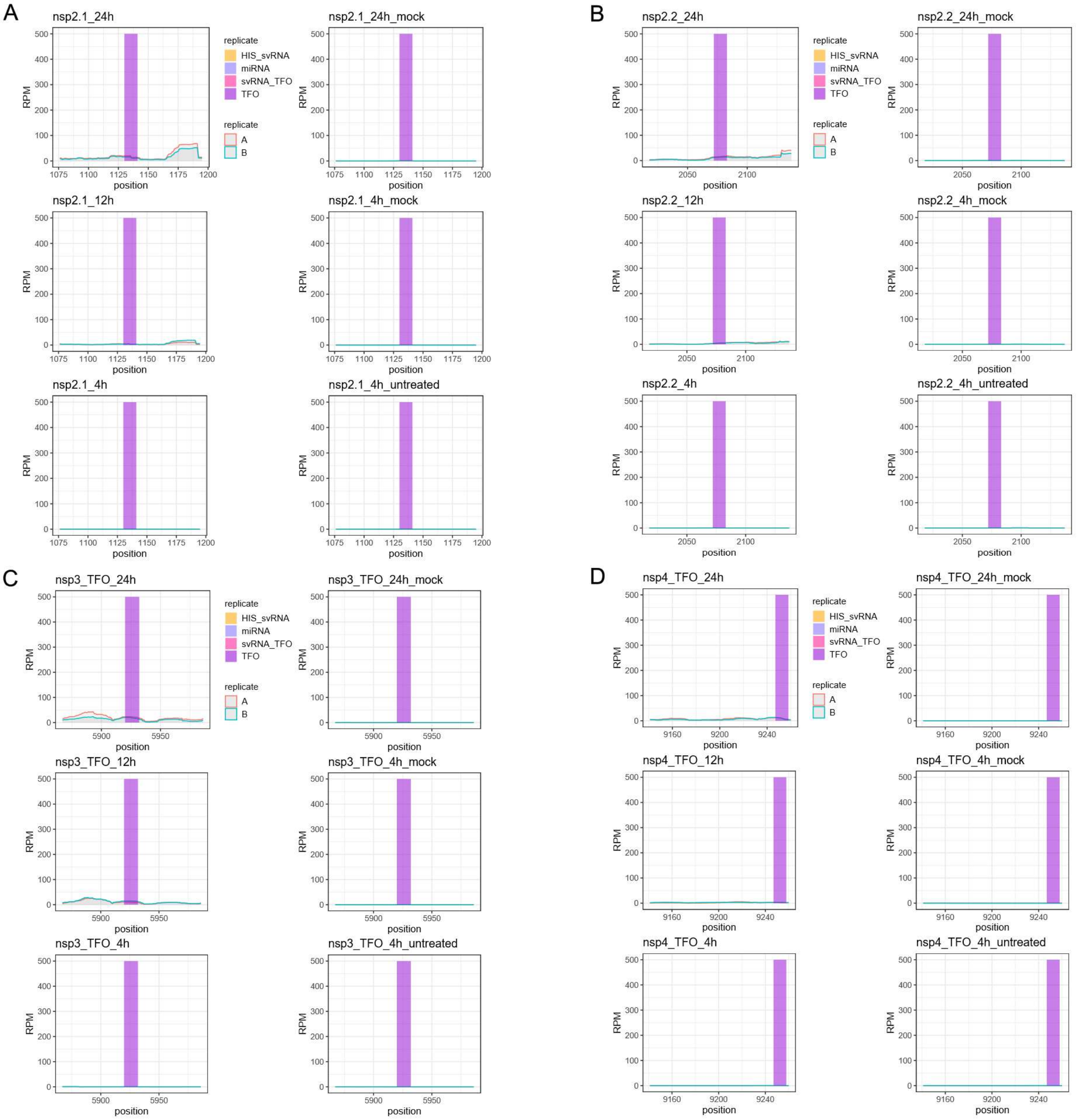

**Supplemental Figure S3. Single-nucleotide coverage of small RNA-sequencing reads (A and B replicate) from SARS-CoV-2-infected Calu-3 cells at 24, 12, and 4 hpi, with mock-infected and untreated as controls [1]**. TFO encoding regions: **A)** portion of the nsp2 ORF region (1,076 – 1,195). **B)** portion of nsp2 ORF region (2,018 – 2,137). **C)** portion of nsp3 ORF region (5,867 – 5,986). **D)** portion of nsp4 ORF region (9,141 – 9,260). The y-axis is presented as the number of reads per million (RPM) and all positions are relative to the Wuhan-Hu-1 genome.

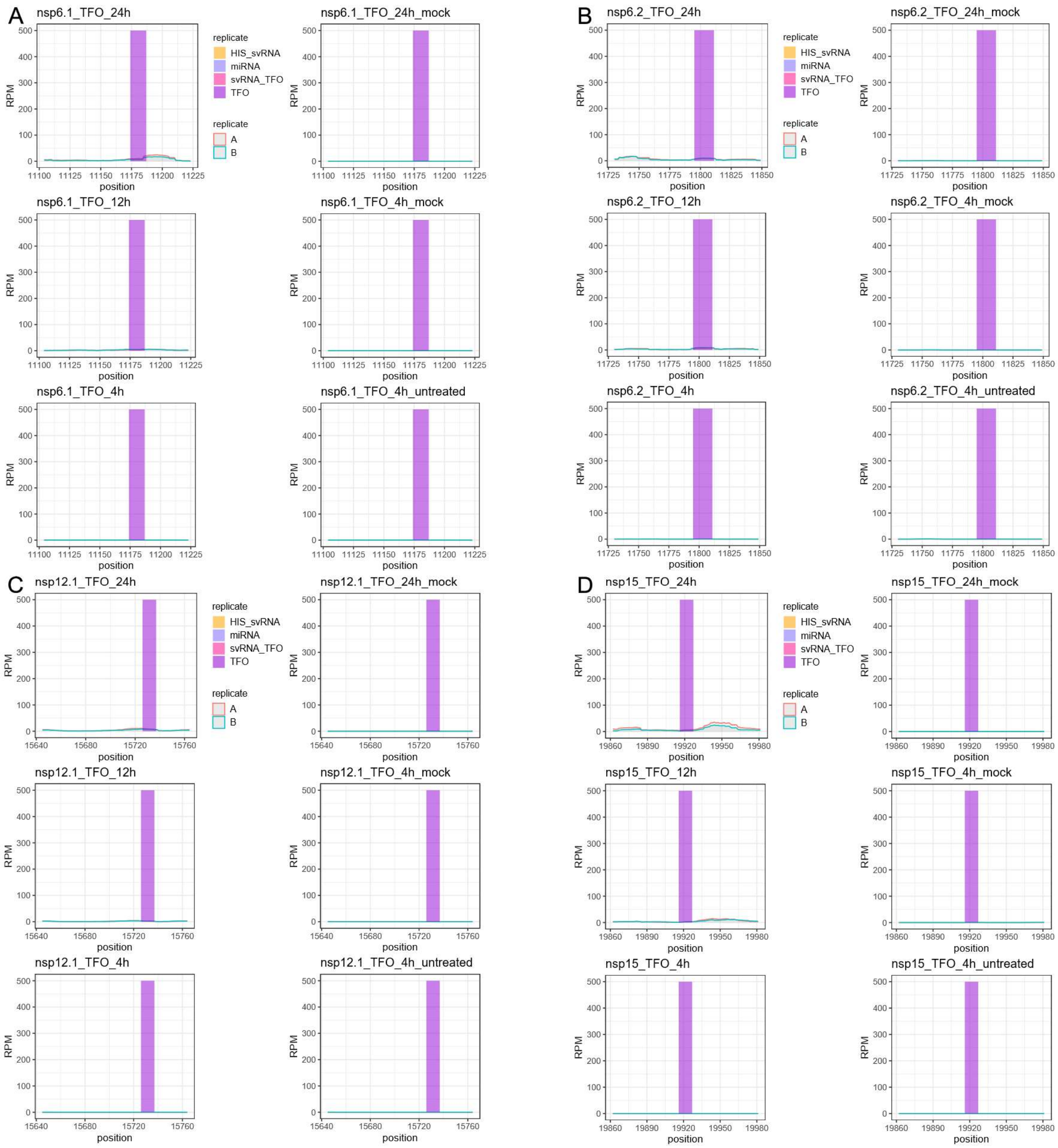

**Supplemental Figure S4. Single-nucleotide coverage of small RNA-sequencing reads (A and B replicate) from SARS-CoV-2-infected Calu-3 cells at 24, 12, and 4 hpi, with mock-infected and untreated as controls** [1]. TFO encoding regions: **A)** portion of the nsp6 ORF region (11,104 – 11,223). **B)** portion of nsp6 ORF region (11,730 – 11,849). **C)** portion of nsp12 ORF region (15,645 – 15,764). **D)** portion of nsp15 ORF region (19,862 – 19,981). The y-axis is presented as the number of reads per million (RPM) and all positions are relative to the Wuhan-Hu-1 genome.

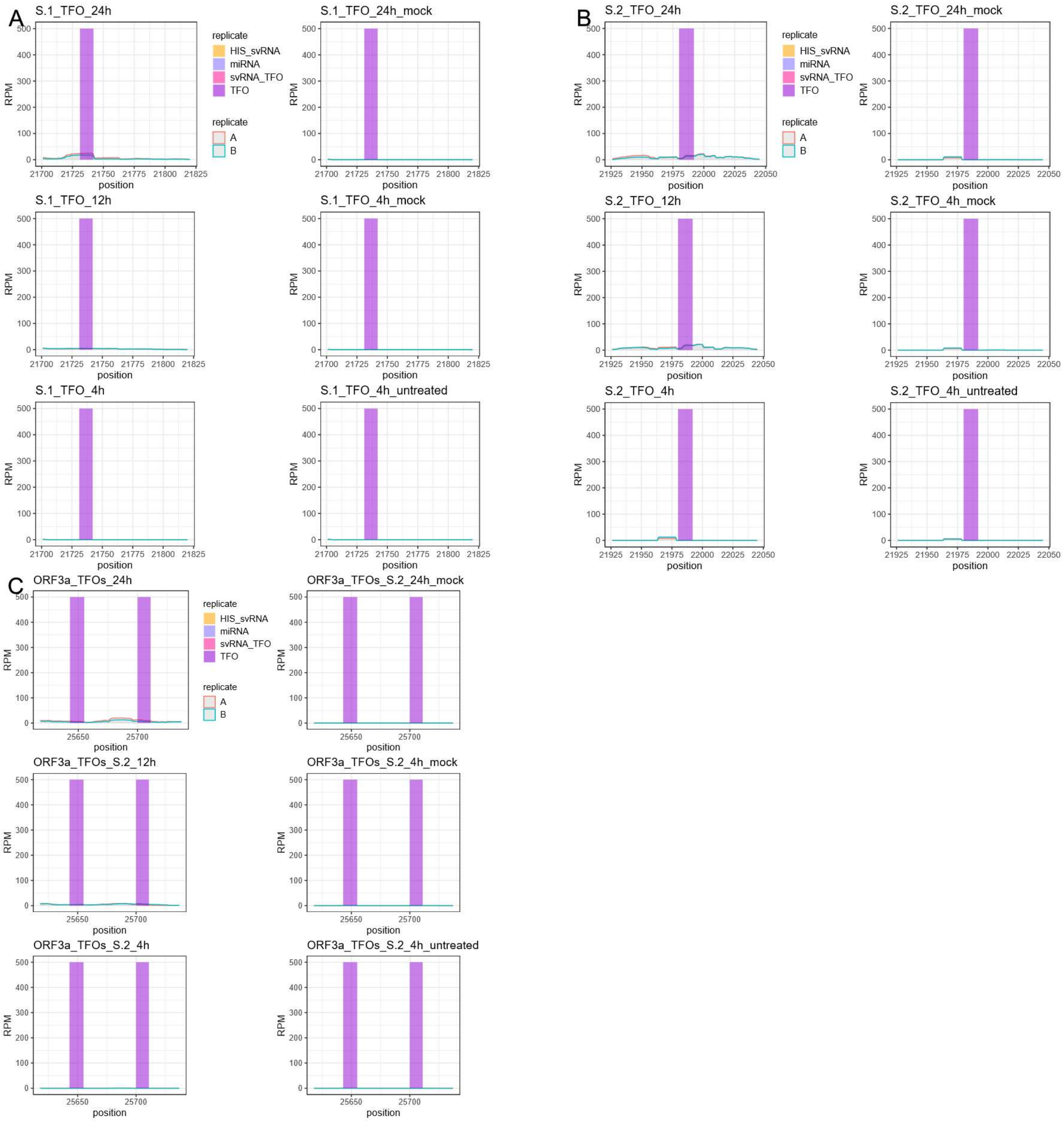

**Supplemental Figure S5. Single-nucleotide coverage of small RNA-sequencing reads (A and B replicate) from SARS-CoV-2-infected Calu-3 cells at 24, 12, and 4 hpi, with mock-infected and untreated as controls [1]. TFO encoding regions: A)** portion of the spike ORF region (21,701 – 21,820). **B)** portion of spike ORF region (21,926 – 22,045). C) portion of ORF3a region (25,618 – 25,737). The y-axis is presented as the number of reads per million (RPM) and all positions are relative to the Wuhan-Hu-1 genome.

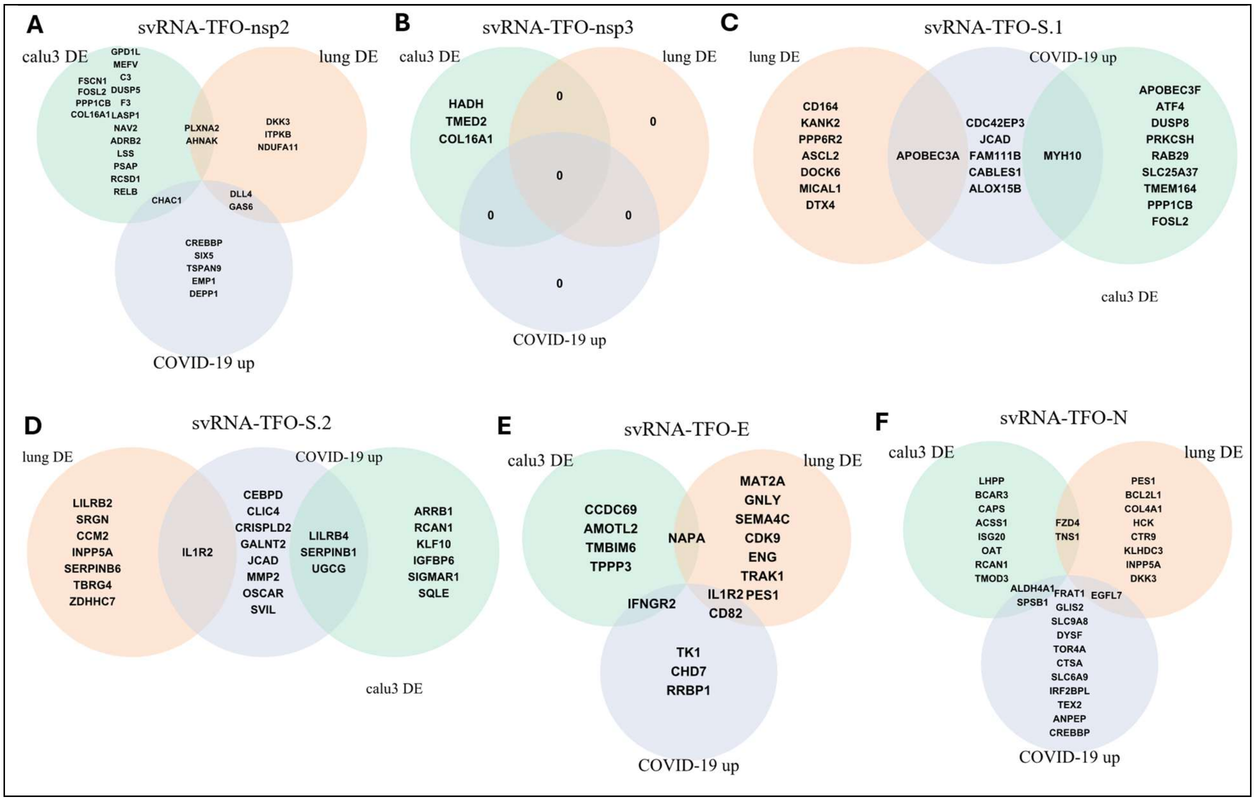

**Supplemental Figure S6. Venn diagrams identifying the most likely human lung enhancer-gene targets of the small viral RNA (svRNA) triplex-forming oligonucleotides (TFOs) of concern. Predicted enhancer-gene targets of A)** svRNA-TFO-nsp2; **B)** svRNA-TFO-nsp3; **C)** svRNA-TFO-S.1; **D)** svRNA-TFO-S.2; **E)** svRNA-TFO-E; and **F)** svRNA-TFO-N. This analysis was based on the topmost differentially expressed (DE) genes in response to SARS-CoV-2 infection of Calu-3 cells, lungs [1,2] and whole blood transcriptome of patients with severe COVID-19 [3].

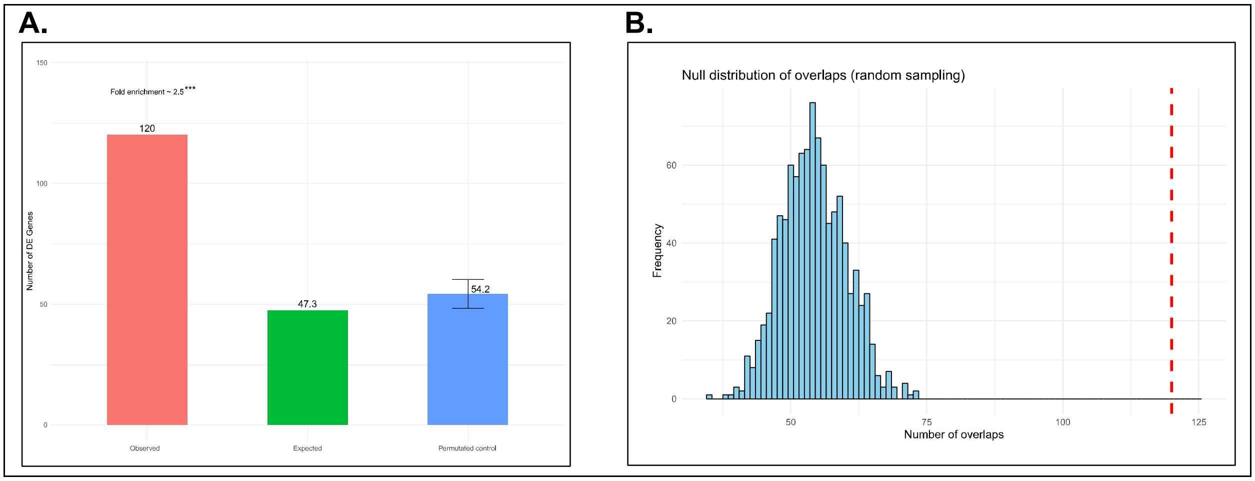

**Supplemental Figure S7. Enrichment analysis of svRNA-TFO gene targets among differentially expressed genes (DEGs) relevant to SARS-CoV-2 infection in Calu-3 cells (GSE148729, GSE147507) and human lung tissues (GSE147507), and upregulated during severe COVID-19 (GSE171110). A)** 120 DEGs were "Observed" to overlap with the 350 svRNA-TFO gene targets identified in this study (Supplemental Figure S6A-F). This represents a significant (***) ~ 2.5-fold enrichment ($p$-value < .001; Fisher's exact test) when compared to the "Expected" number of DEGs based on a background rate of 13.5 %. See Panel B for the distribution of the "Permutated control". **B)** Using the null gene target set, which corresponds to targets of 12 non-svRNA-TFOs detected in this study (see Materials and Methods), 1,000 groups of 350 gene targets were randomly selected to generate the "Permutated control", for which the distribution is shown. This distribution was compared against the 120 DEGs that were "Observed" (represented as the vertical dashed red line), which represent $p$-value < .001 based on a one-sided test $t$-test. See also Supplemental Table S3.

**Supplemental Table S3.** Differentially expressed genes (DEGs) and svRNA-TFO target gene lists and annotations used in the enrichment analysis (see Supplemental Figure S7).

20251229_Supplemental_Table_S3.xlsx

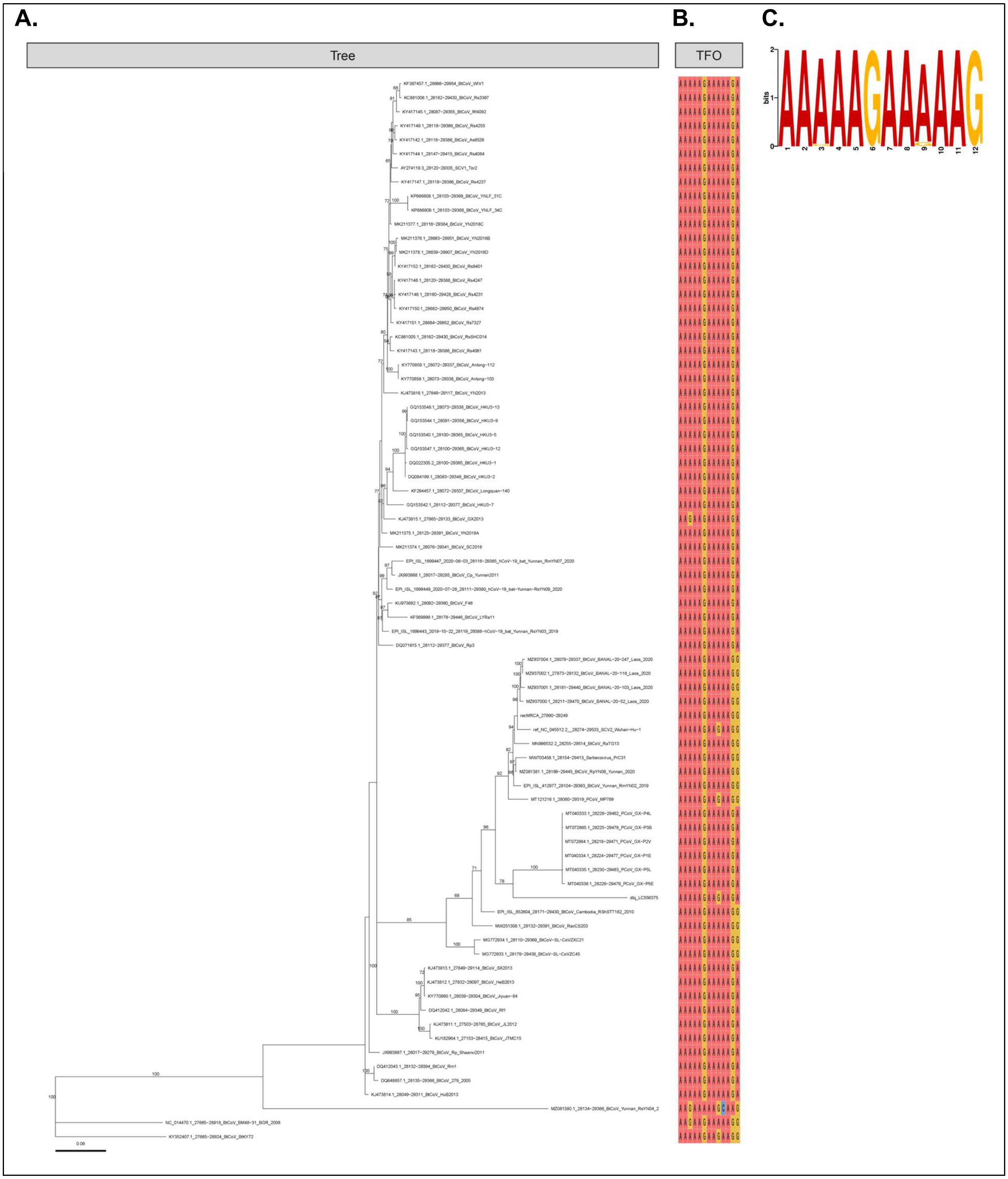

**Supplemental Figure S8. Evolution of Nucleocapsid (N) open reading frame in *Sarbecovirus*. A)** Phylogenetic reconstruction of aligned translated amino acids, with internal node confidence reported for values greater than 65 % out of 10,000 ultrafast bootstraps and re-rooted using Btk72 for visualization purposes. **B)** Sequence alignment segments showing polypurine triplex-forming oligonucleotide (TFO) region. **C)** Motif from 75 TFO-like sequences among *Sarbecovirus*.

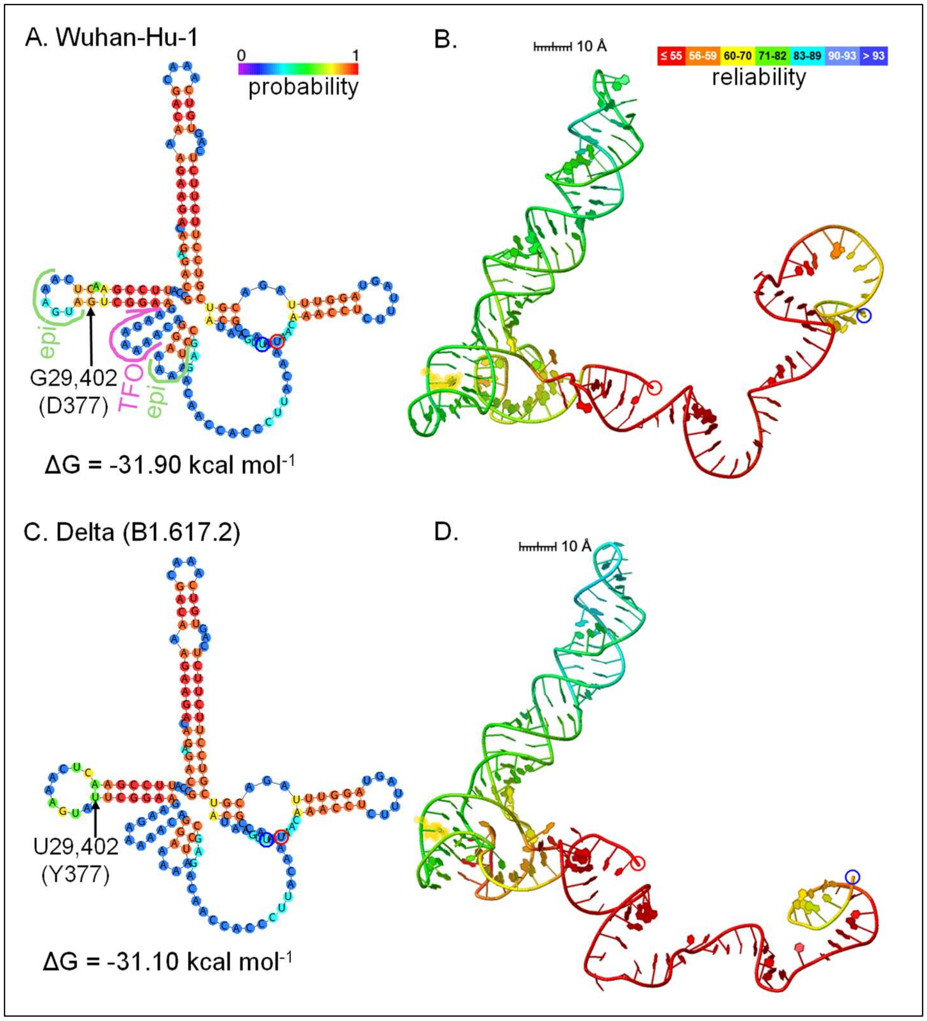

**Supplemental Figure S9. Maximum Free Energy (MFE) of optimal and 3D models for SARS-CoV-2 svRNA-TFO-N progenitor structure (29,343 – 29,492 in Wuhan-Hu-1). A)** MFE of optimal structure for G29402 (D377) variant with base probability indicated by colour using Gibbs free energy (ΔG) and Turner energy model [4] at 37 °C. The AAGAA-like motifs identified in Kim et al. [5] are labelled green as 'epi' and triplex-forming oligo-nucleotides is labelled pink as "TFO". **B)** 3D model for the G29402 (D377) variant with base reliability indicated by colour (see methods). **C)** Same as A, but for U29402 (Y377) variant. **D)** Same as B, but for U29402 (Y377) variant. The 5′ and 3′ ends are highlighted in blue and red, respectively. Yellow highlighting in B and D denotes the G29402U mutation on the 3D models.

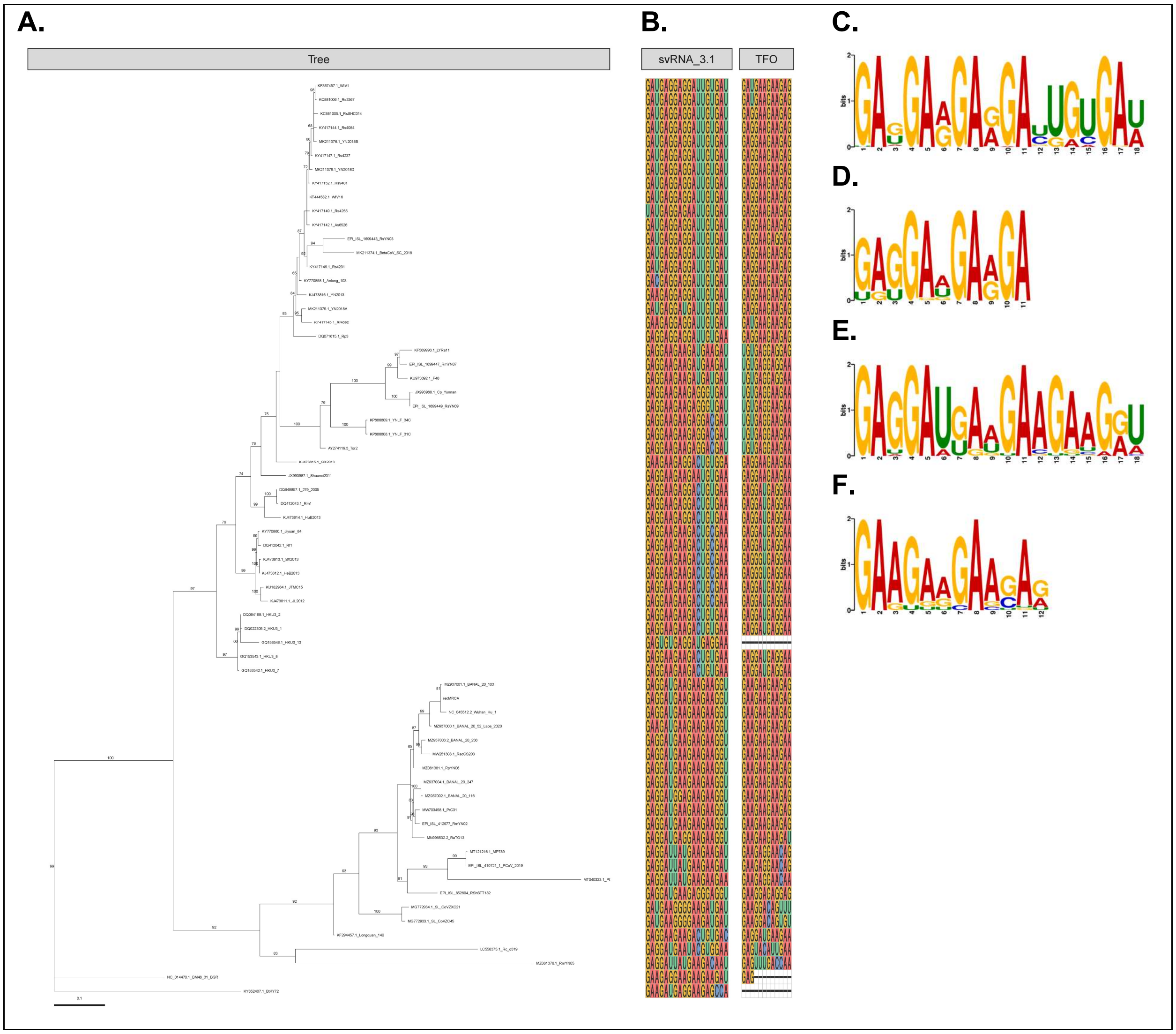

**Supplemental Figure S10. Evolution of the nsp3 open reading frame (recombination region 2 only) in *Sarbecovirus*. A)** Phylogenetic reconstruction of aligned translated amino acids, with internal node confidence reported for values greater than 65 % out of 10,000 ultrafast bootstraps and re-rooted using Btk72 for visualization purposes. **B)** Sequence alignment segments showing svRNA nsp3.1 from SARS-CoV-1 [6] and polypurine triplex-forming oligonucleotide (TFO) regions. **C to F**: Motifs for svRNA nsp3.1 and TFO regions from the SARS-CoV-1- (**C, D**), and SARS-CoV-2-related (**E, F**) sub-groups, respectively.

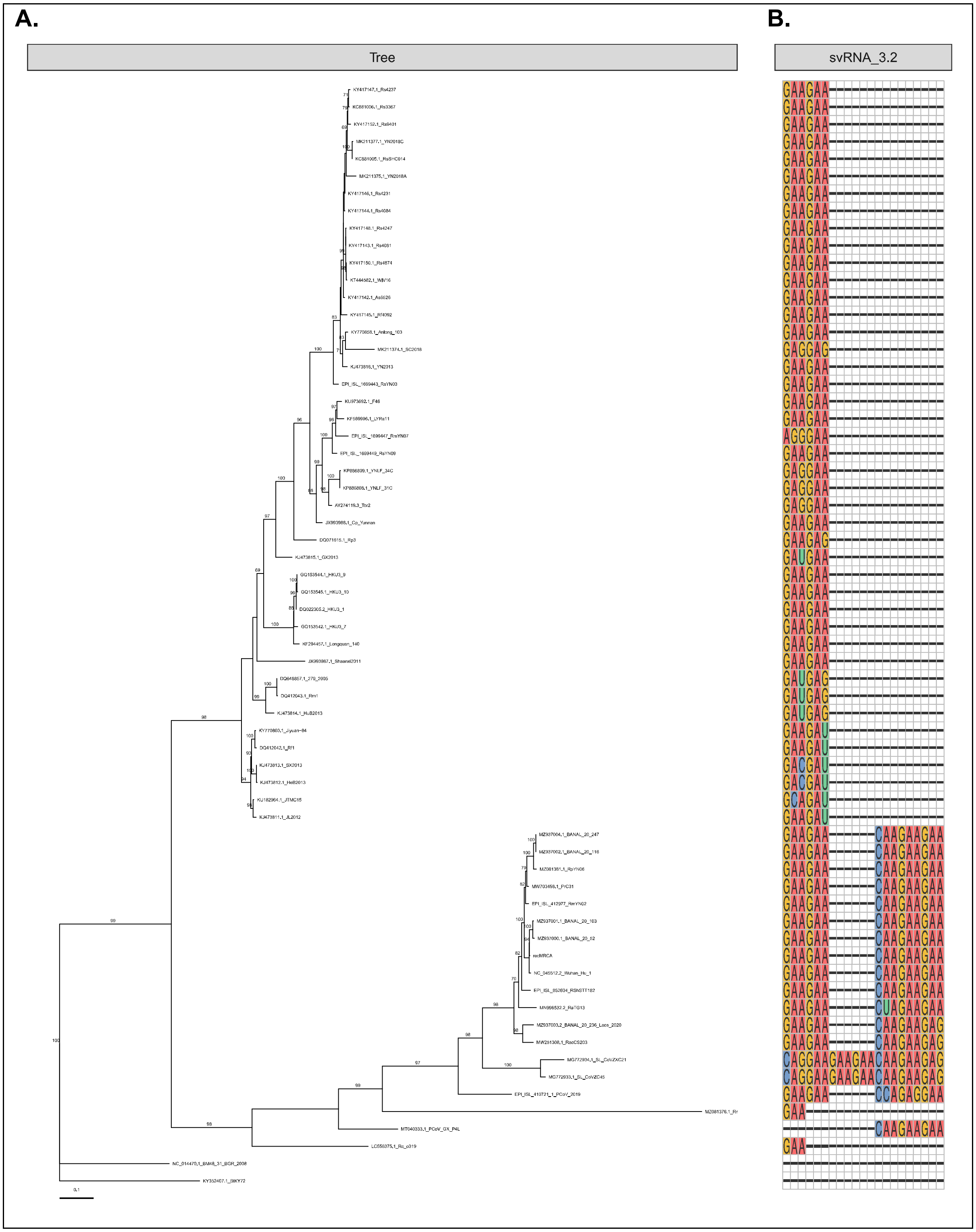

**Supplemental Figure S11. Evolution of the nsp3 open reading frame (recombination region 3 only) in *Sarbecovirus*. A)** Phylogenetic reconstruction of aligned translated amino acids, with internal node confidence reported for values greater than 65 % out of 10,000 ultrafast bootstraps and re-rooted using Btk72 for visualization purposes. **B)** Sequence alignment segments showing presence of an insertion among SARS-CoV-2-related viruses in the svRNA-nsp3.2 region from SARS-CoV-1 [6].

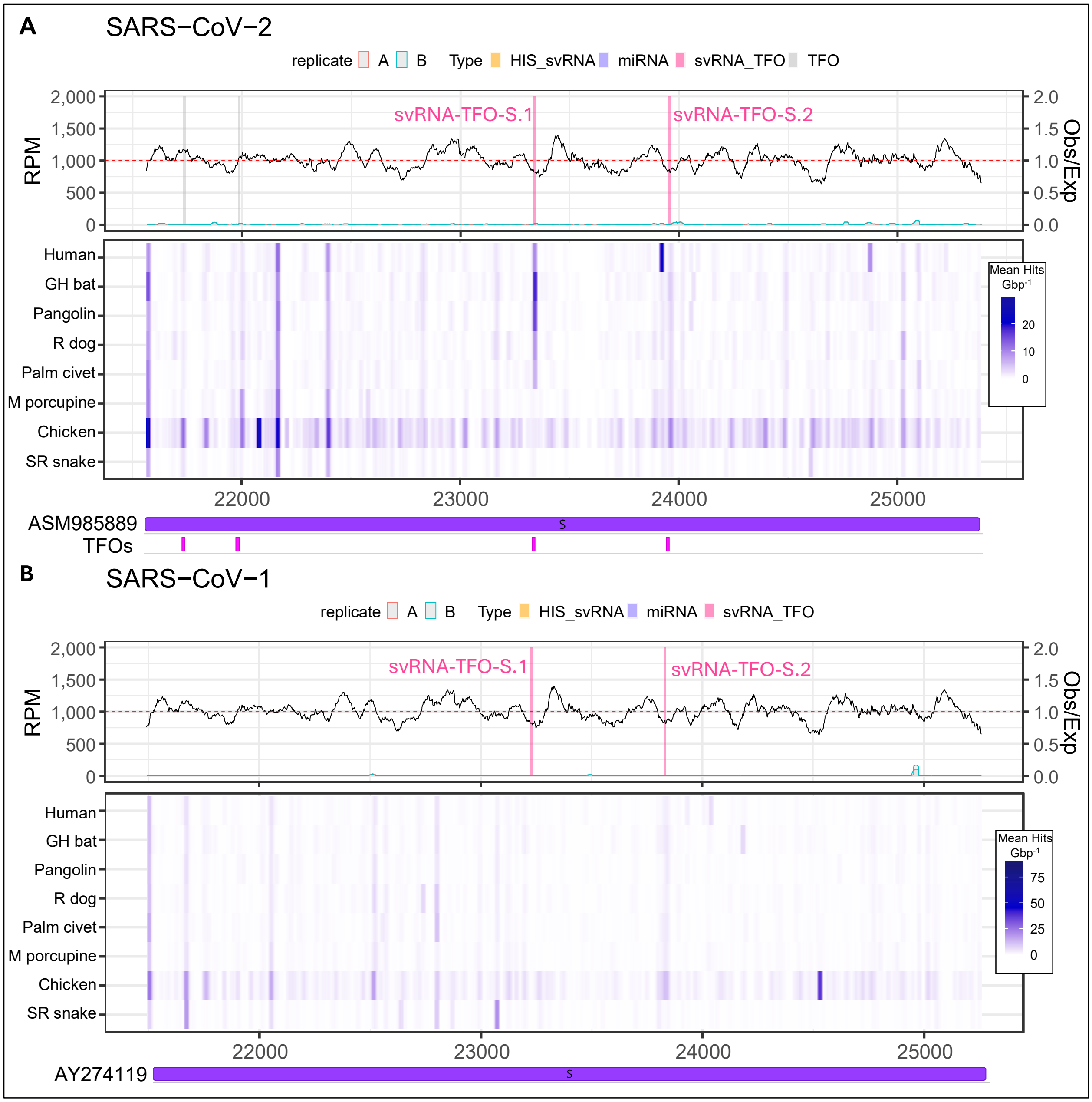

**Supplemental Figure S12. Spike (S) open reading frame region integrating small RNA-sequencing (RNA-seq), synonymous-site conservation (SSC) analysis, triplex-forming oligonucleotide (TFO) predictions, and host genome homology.** Topmost plots, first *y*-axis: reads per million (RPM) coverage small RNA-seq from Calu-3 cells 12 h post-infection [1]. Second *y*-axis: SSC analysis for *Sarbecovirus*, red dashed-line representing an equal ratio of observed (Obs) synonymous mutations compared to the number of expected (Exp). Middle heatmaps: rolling average BLAST hits against various animal genomes (GH bat = greater horseshoe bat; R dog = raccoon dog; M porcupine = Malaysian porcupine; and SR snake = San Diego ring-necked snake) (see Materials and Methods). Bottom feature plots: TFOs predicted against human lung enhancer sequences and svRNA-TFOs of concern identified using a functional genomics pipeline in this study (Supplemental Table S1). **A)** SARS-CoV-2 genetic region 21,563 – 25,384 in Wuhan-Hu-1 reference genome **B)** SARS-CoV-1 genetic region 21,492 – 25,259 in Tor2 reference genome.

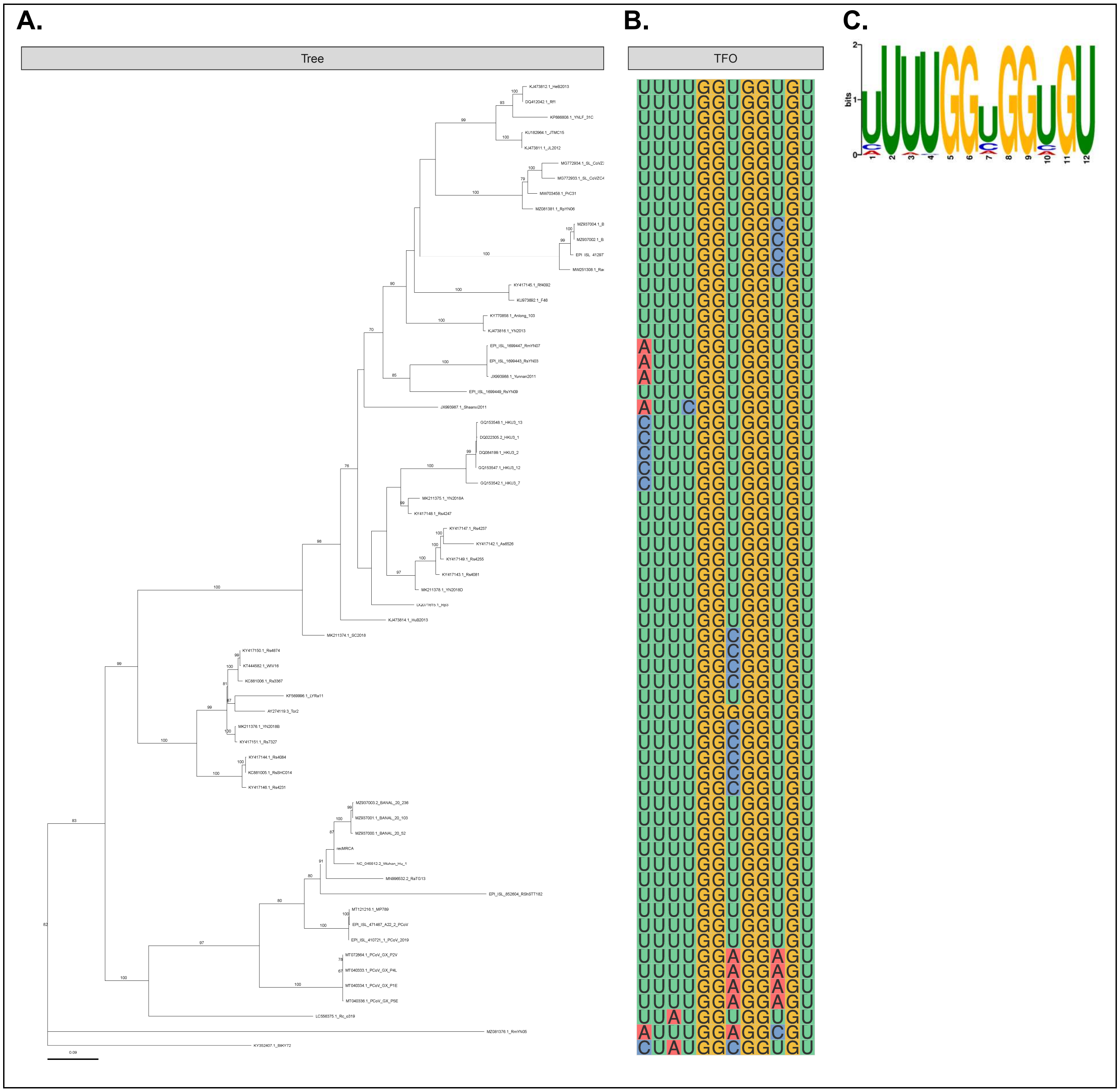

**Supplemental Figure S13. Evolution of Spike (S) open reading frame (recombination region 16 only) in *Sarbecovirus*. A)** Phylogenetic reconstruction of alignment translated amino acids, with internal node confidence reported for values greater than 65 % out of 10,000 ultrafast bootstraps and re-rooted using Btk72 for visualization purposes. **B)** Sequence alignment segments showing purine-pyrimidine triplex-forming oligonucleotide (TFO) associated with svRNA-TFO-S.1 in SARS-CoV-2 (see supplemental Figure S12). **C)** Motif from 64 TFO-like among *Sarbecovirus*.

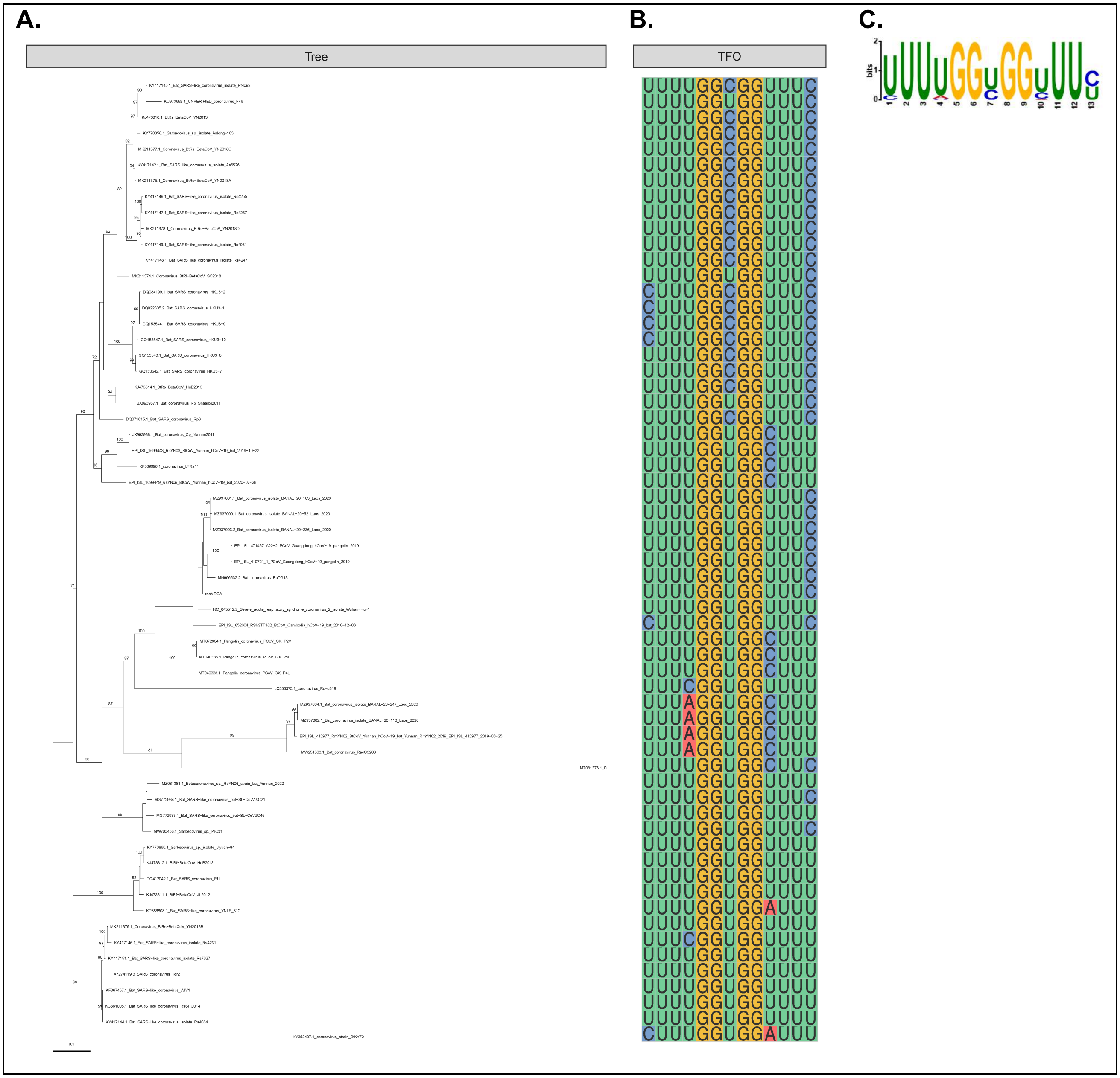

**Supplemental Figure S14. Evolution of Spike (S) open reading frame (recombination region 17 only) in *Sarbecovirus.* A)** Phylogenetic reconstruction of alignment translated amino acids, with internal node confidence reported for values greater than 65 % out of 10,000 ultrafast bootstraps and re-rooted using Btk72 for visualization purposes. **B)** Sequence alignment segments showing purine-pyrimidine triplex-forming oligonucleotide (TFO) associated with svRNA-TFO-S.2 in SARS-CoV-2 (see supplemental Figure S12). **C)** Motif from 61 TFO-like sequence among *Sarbecovirus*.

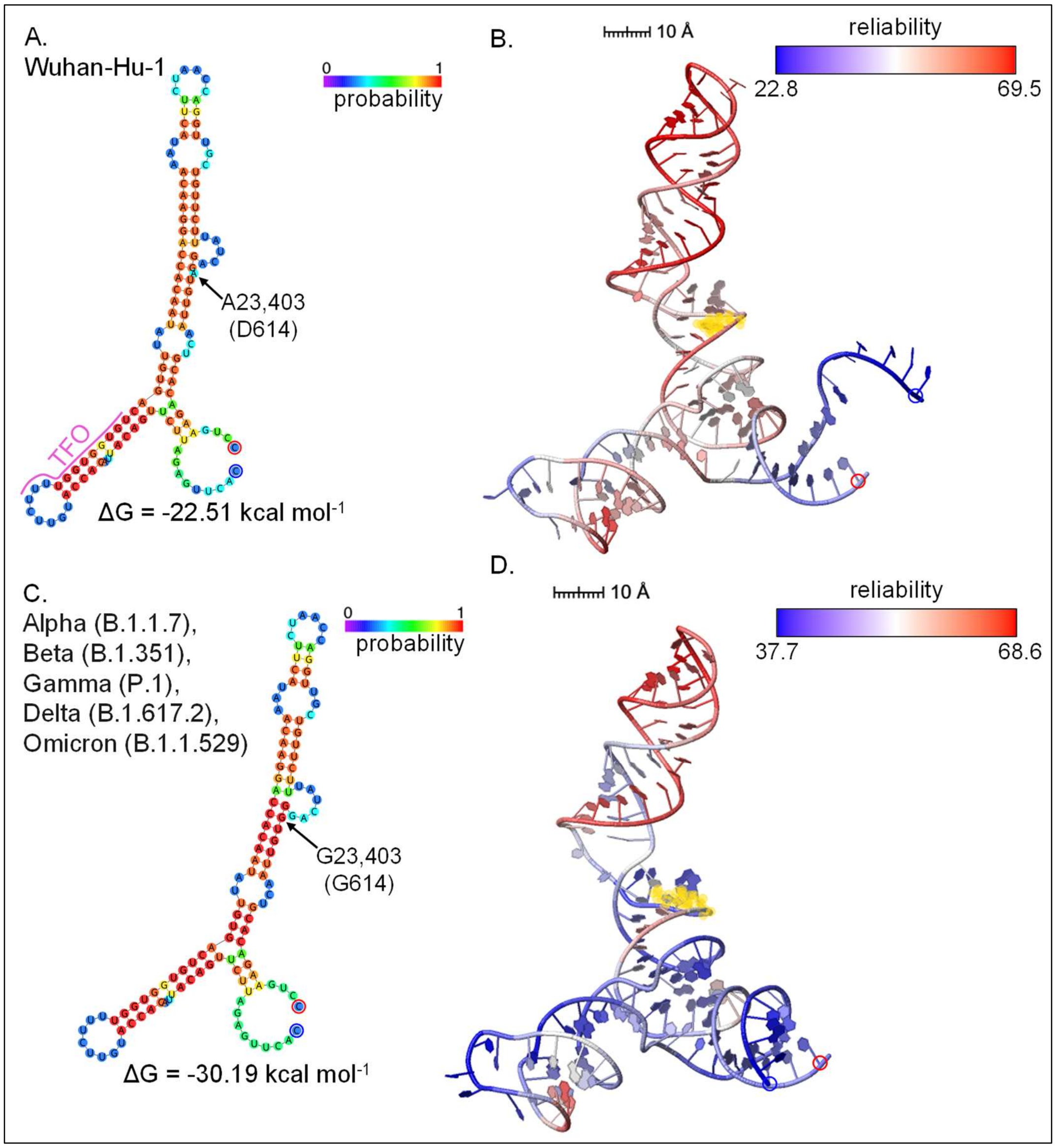

**Supplemental Figure S15. Maximum Free Energy (MFE) of optimal and 3D models for SARS-CoV-2 svRNA-TFO-S.1 progenitor structure (23,304 – 23,423 in Wuhan-Hu-1). A)** MFE of optimal structure for A23403 (D614) variant with base probability indicated by colour using Gibbs free energy (ΔG) and Andronescu energy model [7] at 37 °C. The triplex-forming oligo-nucleotides is labelled pink as "TFO". **B)** 3D model for the A23403 (D614) variant with base reliability indicated by colour. **C)** Same as A, but for G23403 (G614) variant. **D)** Same as

B, but for G23403 (G614) variant. 5′ and 3′ ends are highlighted with blue and red circles, respectively. Yellow highlighting in B and D denotes A23403G (D614G) mutation on 3D models.

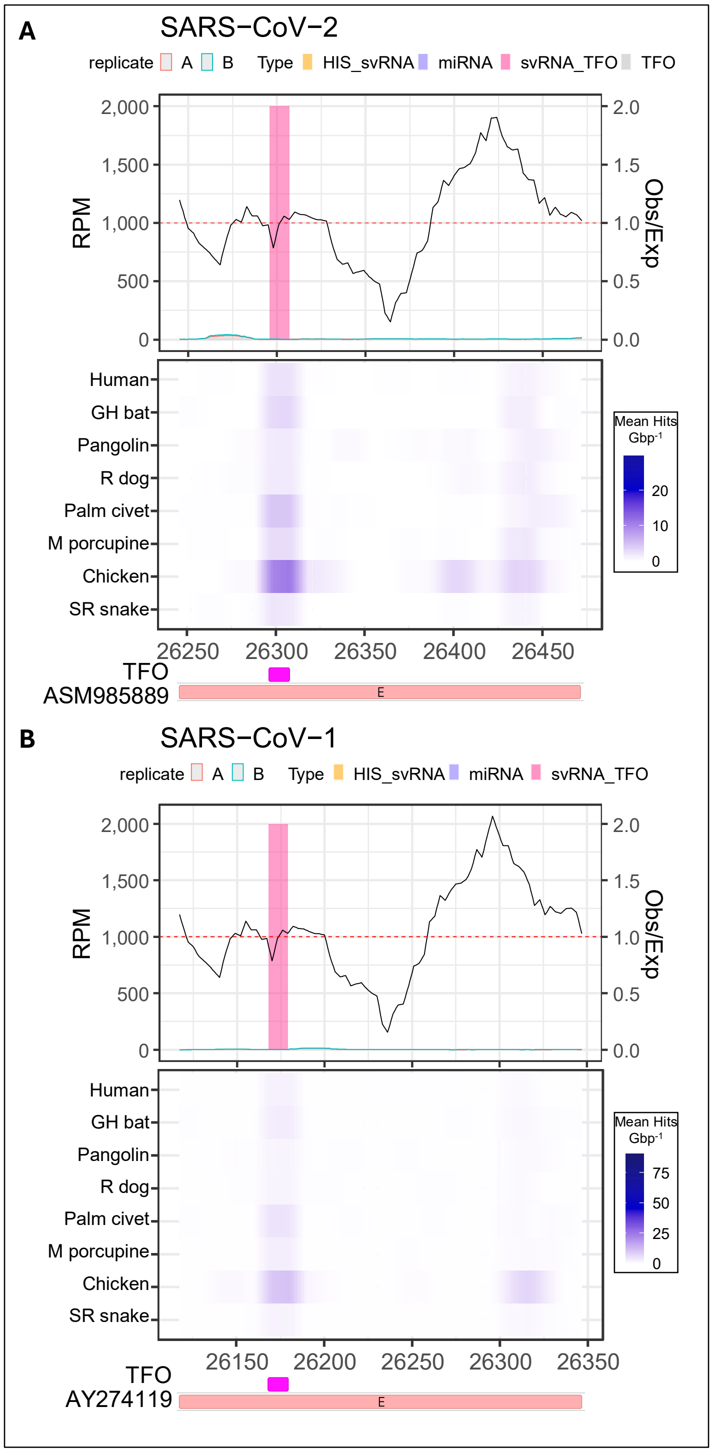

**Supplemental Figure S16. Envelope (E) open reading frame region integrating small RNA-sequencing (RNA-seq), synonymous-site conservation (SSC) analysis, triplex-forming oligonucleotide (TFO) prediction, and host genome homology.** Topmost plots, first *y*-axis: reads per million (RPM) coverage small RNA-seq from Calu-3 cells 12 h post-infection [1]. Second *y*-axis: SSC analysis for *Sarbecovirus*, red dashed-line representing an equal ratio of observed (Obs) synonymous mutations compared to the number of expected (Exp). Middle heatmaps: rolling average BLAST hits against various animal genomes (GH bat = greater horseshoe bat; R dog = raccoon dog; M porcupine = Malaysian porcupine; and SR snake = San

Diego ring-necked snake) (see Materials and Methods). Bottom feature plots: TFO predicted against human lung enhancer sequence and svRNA-TFOs of concern identified using a functional pipeline in this study (supplemental Table S1). **A)** SARS-CoV-2 genetic region 26,245 – 26,472 in the Wuhan-Hu-1 reference genome. **B)** SARS-CoV-1 genetic region 26,117 – 26,347 in the Tor2 reference genome.

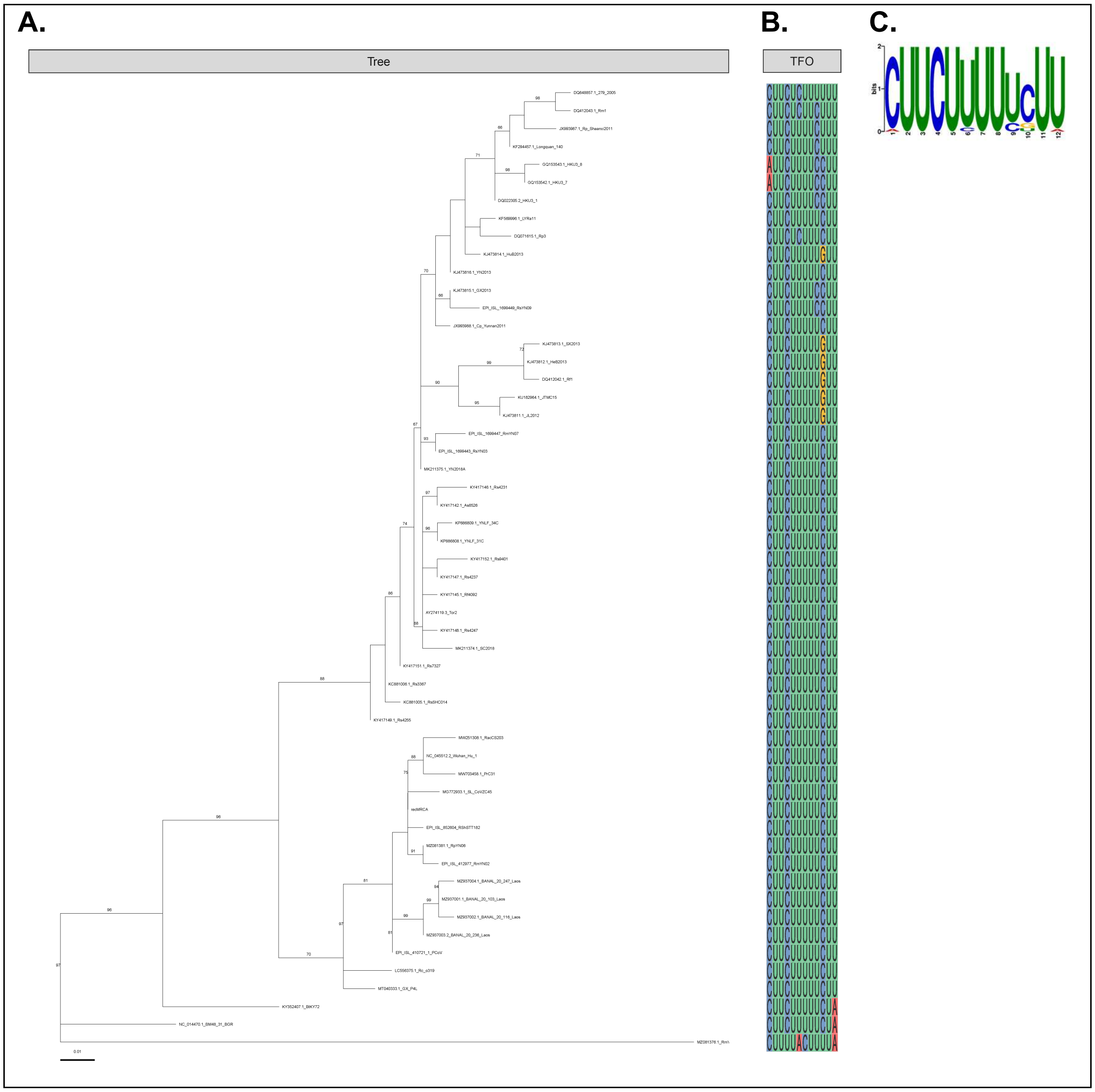

**Supplemental Figure S17. Evolution of Envelope (E) open reading frame (includes recombination regions 19 and 20) in *Sarbecovirus*. A)** Phylogenetic reconstruction of aligned translated amino acids, with internal node confidence reported for values greater than 65 % out of 10,000 ultrafast bootstraps and re-rooted using RmYN05 (MZ081376.1) for visualization purposes. **B)** Sequence alignment segment showing polypyrimidine triplex-forming oligonucleotide (TFO) associated with svRNA-TFO-E in SARS-CoV-2 (see supplemental Figure S16). **C)** Motif from 54 TFO-like sequences among *Sarbecovirus*.

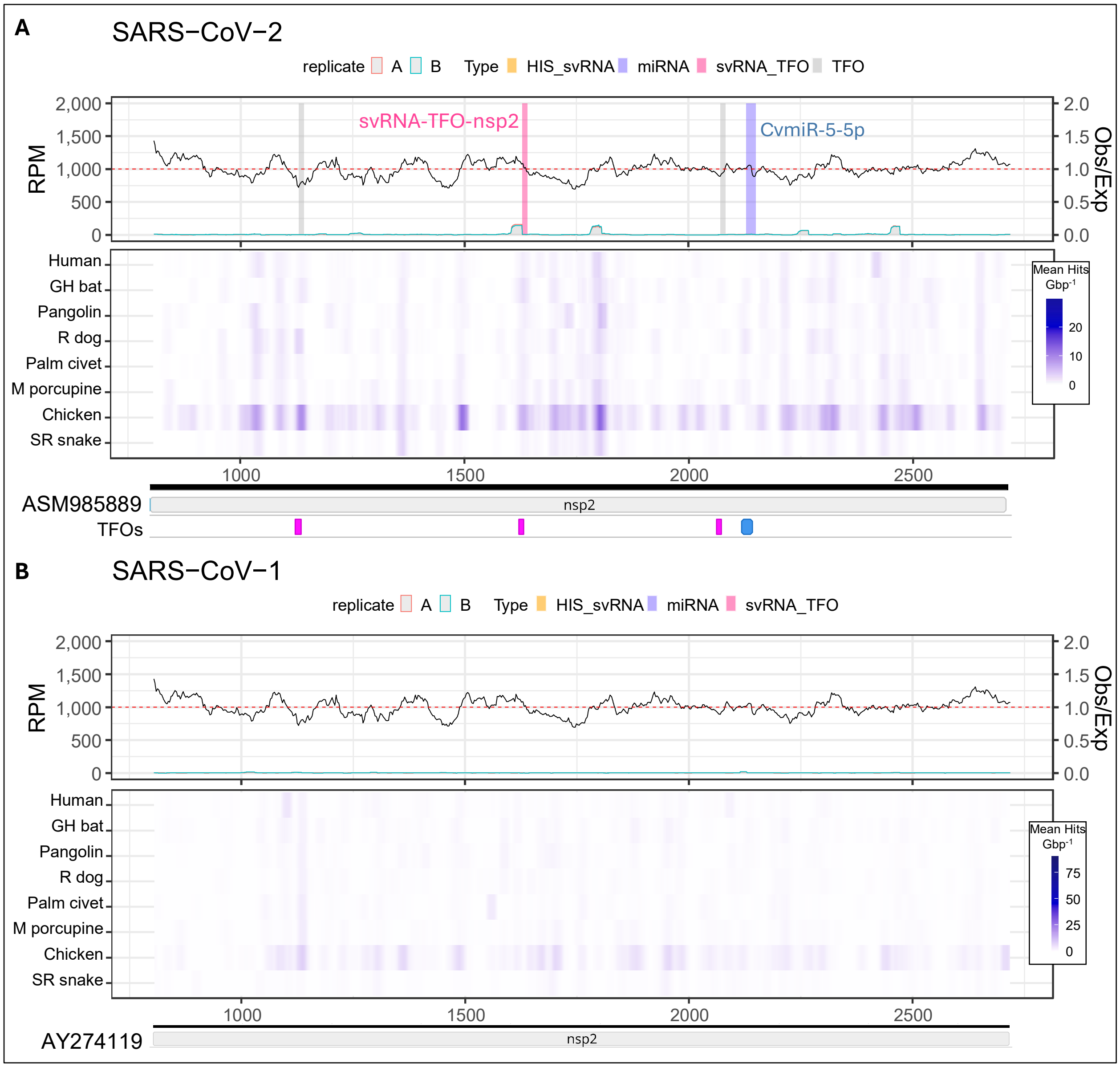

**Supplemental Figure S18. nsp2 region integrating small RNA-sequencing (RNA-seq), synonymous-site conservation (SSC) analysis, microRNA (miRNA) annotations, triplex-forming oligonucleotide (TFO) predictions, and host genome homology.** Topmost plots, first *y*-axis: reads per million (RPM) coverage small RNA-seq from Calu-3 cells 12 h post-infection [1]. Second *y*-axis: SSC analysis for *Sarbecovirus*, red dashed-line representing an equal ratio of observed (Obs) synonymous mutations compared to the number of expected (Exp). Middle heatmaps: rolling average BLAST hits against various animal genomes (GH bat = greater horseshoe bat; R dog = raccoon dog; M porcupine = Malaysian porcupine; and SR snake = San Diego ring-necked snake) (see Materials and Methods). Bottom feature plots: TFOs predicted against human lung enhancer sequences and svRNA-TFOs of concern identified using a functional genomics pipeline in this study (supplemental Table S1); and CvmiR-5-5p was identified by Zhao et al. [8]. **A)** SARS-CoV-2 genetic region 806 – 2,719 in Wuhan Hu-1 reference genome **B)** SARS-CoV-1 genetic region 805 – 2,718 in Tor2 reference genome.

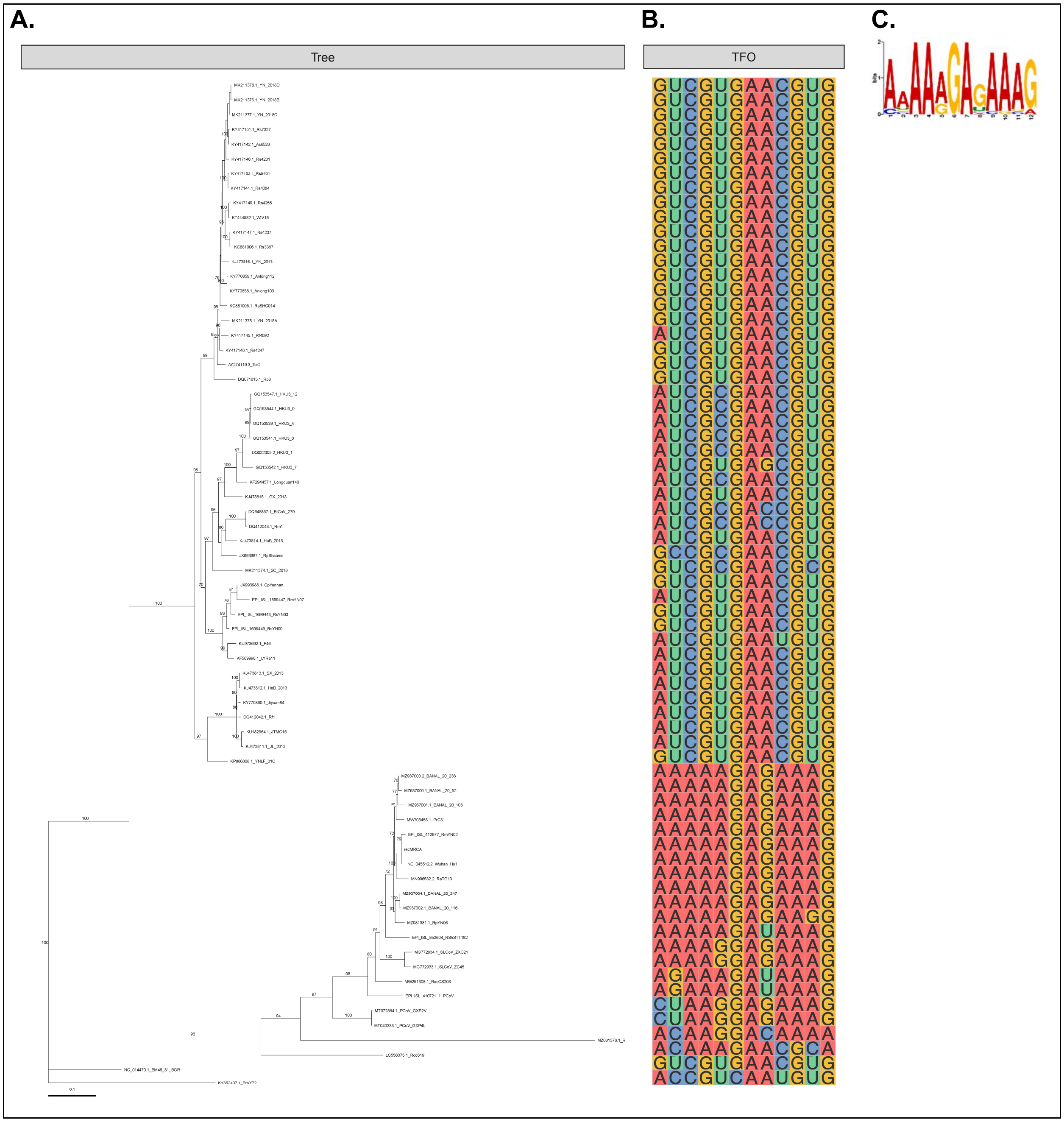

**Supplemental Figure S19. Evolution of partial nsp2 region in *Sarbecovirus*. A)** Phylogenetic reconstruction of aligned translated amino acids, with internal node confidence reported for values greater than 65 % out of 10,000 ultrafast bootstraps and re-rooted using Btk72 for visualization purposes. **B)** Sequence alignment segment showing polypurine triplex-forming oligonucleotide (TFO) associated with svRNA-TFO-nsp2 in SARS-CoV-2 (see supplemental Figure S18). **C)** Motif from 20 aligned TFO-like sequences among the SARS-CoV-2 group.

**Supplemental Table S4.** Sources and accession numbers for SARS-related coronavirus genomes used for phylogenetic reconstruction and synonymous-site conservation analysis for nsp2, nsp3, S-, E-, and N-ORF regions.

20250806_Supplemental_Table_S4.xlsx

**Supplemental Table S5.** Animal host genome resources used in this study

20250806_Supplemental_Table_S5.xlsx
